# RCC1 depletion drives protein transport defects and rupture in micronuclei

**DOI:** 10.1101/2024.09.04.611299

**Authors:** Molly G Zych, Maya Contreras, Manasvita Vashisth, Anna E Mammel, Gavin Ha, Emily M Hatch

## Abstract

Micronuclei (MN) are a commonly used marker of chromosome instability that form when missegregated chromatin recruits its own nuclear envelope (NE) after mitosis. MN frequently rupture, which results in genome instability, upregulation of metastatic genes, and increased immune signaling. MN rupture is linked to NE defects, but the cause of these defects is poorly understood. Previous work from our lab found that chromosome identity correlates with rupture timing for small MN, *i.e.* MN containing a short chromosome, with more euchromatic chromosomes forming more stable MN with fewer nuclear lamina gaps. Here we demonstrate that histone methylation promotes rupture and nuclear lamina defects in small MN. This correlates with increased MN size, and we go on to find that all MN have a constitutive nuclear export defect that drives MN growth and nuclear lamina gap expansion, making the MN susceptible to rupture. We demonstrate that these export defects arise from decreased RCC1 levels in MN and that additional loss of RCC1 caused by low histone methylation in small euchromatic MN results in additional import defects that suppress nuclear lamina gaps and MN rupture. Through analysis of mutational signatures associated with early and late rupturing chromosomes in the Pan-Cancer Analysis of Whole Genomes (PCAWG) dataset, we identify an enrichment of APOBEC and DNA polymerase E hypermutation signatures in chromothripsis events on early and mid rupturing chromosomes, respectively, suggesting that MN rupture timing could determine the landscape of structural variation in chromothripsis. Our study defines a new model of MN rupture where increased MN growth, caused by defects in protein export, drives gaps in nuclear lamina organization that make the MN susceptible to membrane rupture with long-lasting effects on genome architecture.

## Introduction

Micronuclei (MN) form when missegregated chromatin recruits its own nuclear envelope (NE) during mitotic exit and can contain chromatin fragments or one or a few whole chromosomes. MN are used as markers of exposure to carcinogens, are frequent in many tumors, and occur at elevated rates in genetic diseases, including auto-immune disorders and laminopathies^1,2^. Similar to nuclei, the micronuclear envelope compartmentalizes chromatin, however many nuclear functions are defective in MN, including chromatin modifications, transcription, replication, and nuclear import^3^. Because of these defects, micronucleation frequently causes functional aneuploidy, where a chromosome is diploid but one copy is not expressed, and to DNA damage from erroneous transcription and replication^4,5^. MN are extremely unstable; in many cultured cell lines at least 50% of MN undergo membrane rupture and lose compartmentalization during interphase with the proportion increases with increasing cell cycle duration^6^. Rupture exposes the chromatin to cytosolic enzymes and interrupts DNA replication, leading to massive DNA damage, aneuploidy, and extensive highly localized genome changes, including chromothripsis and kataegis^7^. MN rupture also activates the cGAS-STING innate immune pathway, which can trigger cancer-associated invasion and inflammatory signaling^8–10^.

A current model for MN rupture is centered on defects in NE assembly, including decreased density of nuclear pore complexes (NPCs) and gaps in the nuclear lamina that become the site of membrane rupture^6^. Small MN containing shorter chromosomes deficient in the recruitment of “non-core” NE proteins during mitotic exit, including B-type lamins, membrane binding nucleoporins (Nups), and LBR^11,12^. These defects are proposed to arise from high microtubule density or elevated aurora B activity during chromatin missegregation^11,13–15^ and sustained by reduced NPC assembly and protein import^16^. An alternate model proposes that high membrane curvature in small MN inhibits lamin B meshwork assembly^17,18^. Impaired nuclear import is also proposed to underlie reductions of key replication, transcription, and histone modification proteins in MN, leading to their dysfunction^19,20^. However, whether reduced NPC density is sufficient to cause all the reported defects in protein accumulation remains unclear.

We previously found that MN formed around chromosomes with low densities of lamina associated heterochromatin domains (LADs) have improved nuclear lamina organization despite limited recruitment of lamin B1 and NPCs^12^, suggesting that another factor can drive membrane rupture in MN. In this study we demonstrate that histone methylation increases nuclear lamina gap appearance and MN rupture via a mechanism linked to MN growth. We identify reduced nuclear export rates as a major driver of MN growth and nuclear lamina disruption in MN and find that reduced recruitment of the RanGEF RCC1 to MN is the basis of this defect. We demonstrate that RCC1 is further reduced on euchromatic MN due to lower levels of histone methylation, leading to an additional import defect that suppress export cargo accumulation and MN rupture. In addition, analysis of cancer genome datasets, identifies an enrichment of distinct hypermutation signatures on putative early and mid-rupturing chromosome MN with chromothripsis, suggesting that MN rupture timing can have long-lasting impacts on genome structure that could alter cancer progression and therapy efficacy.

## Results

### Heterochromatin decreases MN stability by disrupting micronuclear lamina organization

Previously, we identified a rescue of nuclear lamina gaps in MN containing human chromosome 19 (chr 19), leading to a delay in rupture compared to similarly sized MN formed around the more heterochromatin-rich chromosome 18 (chr 18) (Fig. 1A). These differences in the timing of nuclear lamina gap appearance were not correlated with differences in NPC density or lamin B1 recruitment, as both were decreased to similar levels in these MN compared to larger ones^12^. To identify the mechanism protecting nuclear lamina organization in chr 19 MN, we first asked which euchromatin-associated characteristics were altered between intact chr 18 and chr 19 MN. Single chromosome MN were induced in hTERT RPE-1 cells (RPE1) by adding an Mps1 inhibitor (Mps1i) to cells previously synchronized in G1 by incubation in the Cdk4/6 inhibitor PD-0332991 isethionate for 24 h (Fig. 1F). Cell synchronization ensures that we only analyze MN during the first cell cycle and that differences in MN rupture timing can be robustly analyzed. Single chr 18 and 19 MN were identified by immunofluorescence (IF)-DNA FISH using a centromere marker to find MN with only 1 chromosome and a chromosome paint or enumeration probe to determine the content. Intact MN were identified by the presence of H3K27Ac labeling or the lack of LBR accumulation^12,21^ (Fig. S1A, B). Cells were fixed in late G1 after Mps1i addition and labeled with representative markers of heterochromatin (H3K27me3 and H3K9me2), euchromatin (H3K27ac), and transcription (RNA PolII pSer2). Consistent with previous results^19,20^, H3K9me2 was enriched and active RNA PolII substantially depleted in intact MN compared to nuclei (Fig. S1E-H). Comparing chr 18 and 19 MN, we found enrichment of H3K27me3, a marker of facultative heterochromatin, in chr 18 MN (Fig. 1B-D) and an unexpected enrichment of active RNA PolII in chr 18 MN (Fig. 1E). Overall levels of RNA PolII pSer2 were massively reduced in both chr 18 and 19 MN compared to nuclei (Fig. S1D), though, suggesting that transcription is not contributing to MN stability for either chromosome.

**Figure 1:**
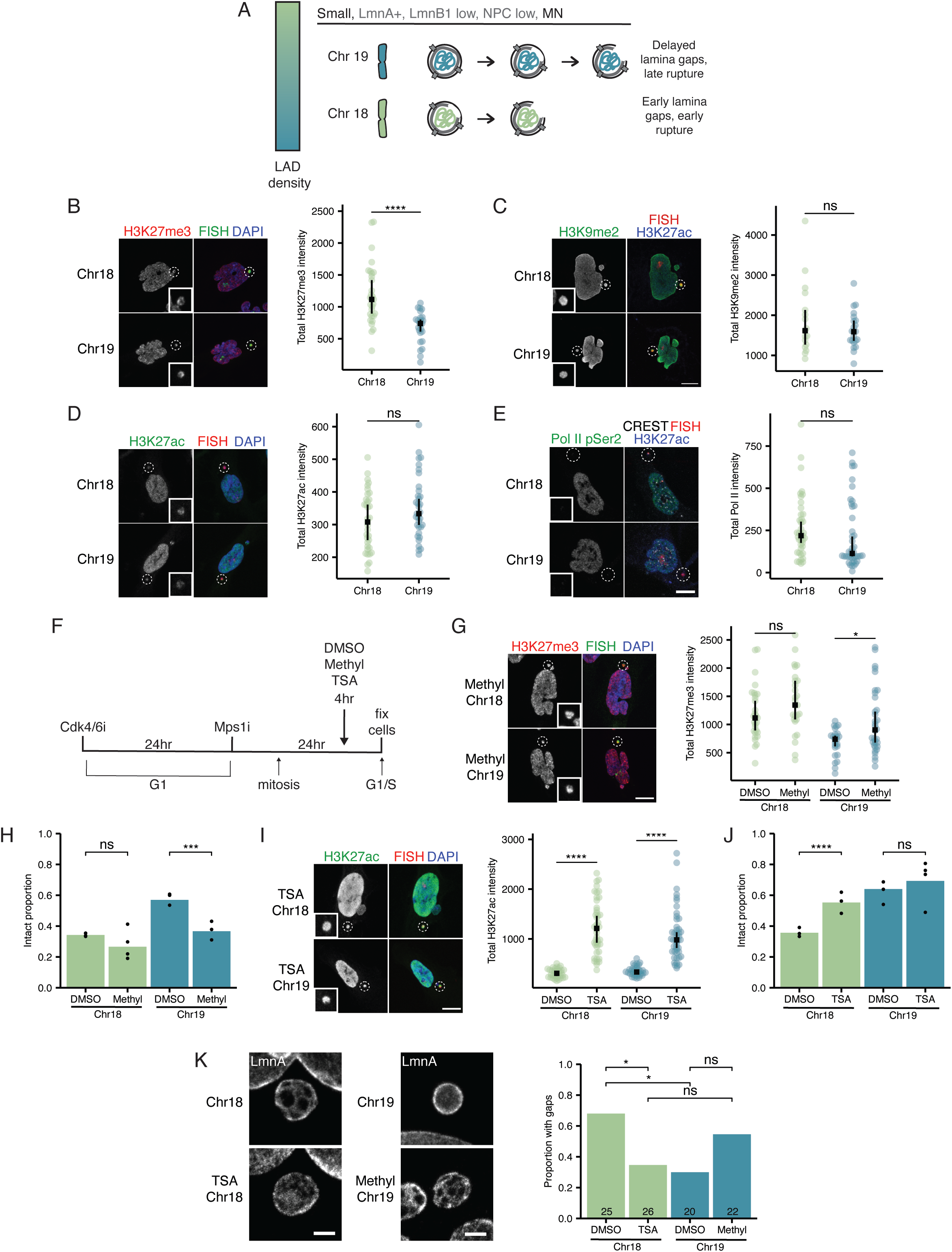
Heterochromatin decreases MN stability by disrupting micronuclear lamina organization. **A.** Summary of the influence of chromosome size and gene density on lamina gap formation and MN rupture timing (Mammel et al). **B-E.** Maximum intensity projections of H3K27me3 (B), H3K9me2 (C), H3K27ac (D), and Pol II pSer2 (E) in intact single chromosome chr 18 and 19 MN in RPE1 cells 24hrs post BAY addition. Quantification H3K27me3, H3K9me2, H3K27ac, and RNA PolII pSer2 from sum projections to measure total intensity. Wilcox ran sum test, N=3, H3K27me3 n=(30,25), H3K9me2 n=(17,22), H3K27ac n=(74,65), and Pol II pSer2 n=(16, 15). **F.** Experimental outline of 4hr methylstat and TSA drug treatments. **G.** Maximum intensity projections of H3K27me3 in intact single chromosome chr 18 and 19 MN in RPE1 cells after 4hr treatment with 5uM methylstat. Wilcoxon rank sum test, N=4, n=(30, 27, 25, 34) **H.** MN stability for 24hr post BAY release single chromosomes chr 18 and 19 MN after 4hr DMSO or 5uM methylstat treatment. Barnards test, N=(4,4,3,3), n=(216,338,165,199). **I.** Maximum intensity projections of H3K27ac in intact single chromosome chr 18 and 19 MN in RPE1 cells after 4hr treatment with 100nM TSA. Scale bars=10um. Quantification H3K27ac from sum projection to measure total intensity. Wilcox ran sum test, N=3, n=(74,65,90,53) **J.** MN stability for 24hr post BAY release single chromosomes chr 18 and 19 MN after 4hr DMSO or 100nM TSA treatment. Barnards test, N=(3,3,4,4), n=(213, 188, 203,112). Scale bars = 10um. **K.** Max projections of the top surface of intact MN containing single chromosome 18 or 19 MN in RPE1 cells. Example of MN without gaps for DMSO chr 19 and TSA chr 18 and example of MN with gaps for DMSO chr 18 and methylstat chr 19. Quantification of nuclear lamina organization in MN for each condition. Barnards test, N=3, n=(25, 26, 20, 22). Scale bar = 1um. For all graphs, ns p>0.05, * p<0.05, ** p<0.01, *** p<0.001, **** p<0.0001. For all dotplots in the article median of all replicates (black square) is shown with 95% confidence interval (black line) and individual measurements (colored circles). For all bar graphs in the article, individual experiments are represented as points and pooled replicate proportions are represented as bars. For all sample sizes, N = number of experimental replicates, n = total number of objects from all replicates.

Based on these results, we asked whether altering histone modifications was sufficient to change MN stability. Addition of either a histone demethylase inhibitor, methylstat, or a histone deacetylase inhibitor, trichostatin A (TSA), to post-mitotic cells for 4 h (Fig. 1F) was sufficient to alter histone modifications in nuclei and bulk MN (Fig. S1E-G) and to avoid off-target effects on chromosome missegregation^22,23^. Methylstat significantly increased H3K27me3 intensity on chr 19 MN (Fig. 1G) and this was correlated with a reduction in MN stability (Fig. 1H). A similar, but less significant trend was observed for chr 18 MN. In contrast, TSA increased H3K27Ac intensity on both chr 18 and 19 MN (Fig. 1I) and was correlated with increased stability of chr 18 MN (Fig. 1J). Interestingly, analysis of bulk MN integrity after TSA and methylstat treatment showed a different effect, with both treatments increasing the proportion of intact MN (Fig. S1I, J). Together, these results suggest that histone methylation, and specifically heterochromatin-associated marks, contributes to the rupture of small MN and that rupture of large MN may be regulated by a distinct mechanism^11,12,24^.

We previously found that nuclear lamina gaps are required for MN rupture and are larger and more frequent in small MN with high LAD densities, *e.g.* chr 18 MN^12,25^. To determine whether histone modifications contributed to MN lamina organization, we used super resolution microscopy of lamin A to ask whether reversing histone modifications in chr 18 and 19 MN altered nuclear lamina gap frequency. Lamin A is recruited to similar levels across all MN, making it an ideal marker for overall nuclear lamina organization^12^. In control cells, chr 18 MN had a higher frequency of lamina gaps compared to chr 19 MN, as expected, and treatment with TSA (chr 18) and methylstat (chr 19) was sufficient to reverse this relationship (Fig. 1K). These data suggest that histone methylation or associated changes in DNA structure drive defects in nuclear lamina organization and accelerate MN rupture.

### MN growth is required for nuclear lamina gaps and MN rupture

Peripheral heterochromatin is a critical regulator of nuclear responses to mechanical force^26–30^. Thus, we asked whether chr 18 and 19 MN differed in the presence or extent of LADs. Analysis of the localization and intensity of the LAD marker H3K9me2 found no difference between chr 18 and 19 MN and little to no effect of methylstat or TSA on this domain (Fig. S2A, B), indicating that peripheral heterochromatin likely does not contribute to MN stability differences.

During our analysis of MN characteristics after methylstat and TSA treatment, we noticed that chr 19 MN incubated in methylstat, which increased nuclear lamina gaps and decreased intact MN, were significantly larger than control chr 19 MN (Fig. 2A). This change was not observed for chr 18 MN or bulk MN (Fig. 2A, Fig. S2C), which did not show decreased integrity after methylstat incubation. We observed a similar correlation between MN size and lamina gap appearance for other single chromosome MN, where MN with lamina gaps were larger on average than ones without (Fig. 2B). To determine whether MN growth was correlated with nuclear lamina gap appearance, we performed live cell imaging of U2OS cells expressing GFP-NLS and mCherry-lamin A. Analysis of MN and lamina gap area found a strong correlation between MN area and lamina gap size over time (Fig. 2C), suggesting a functional connection between these properties.

**Figure 2:**
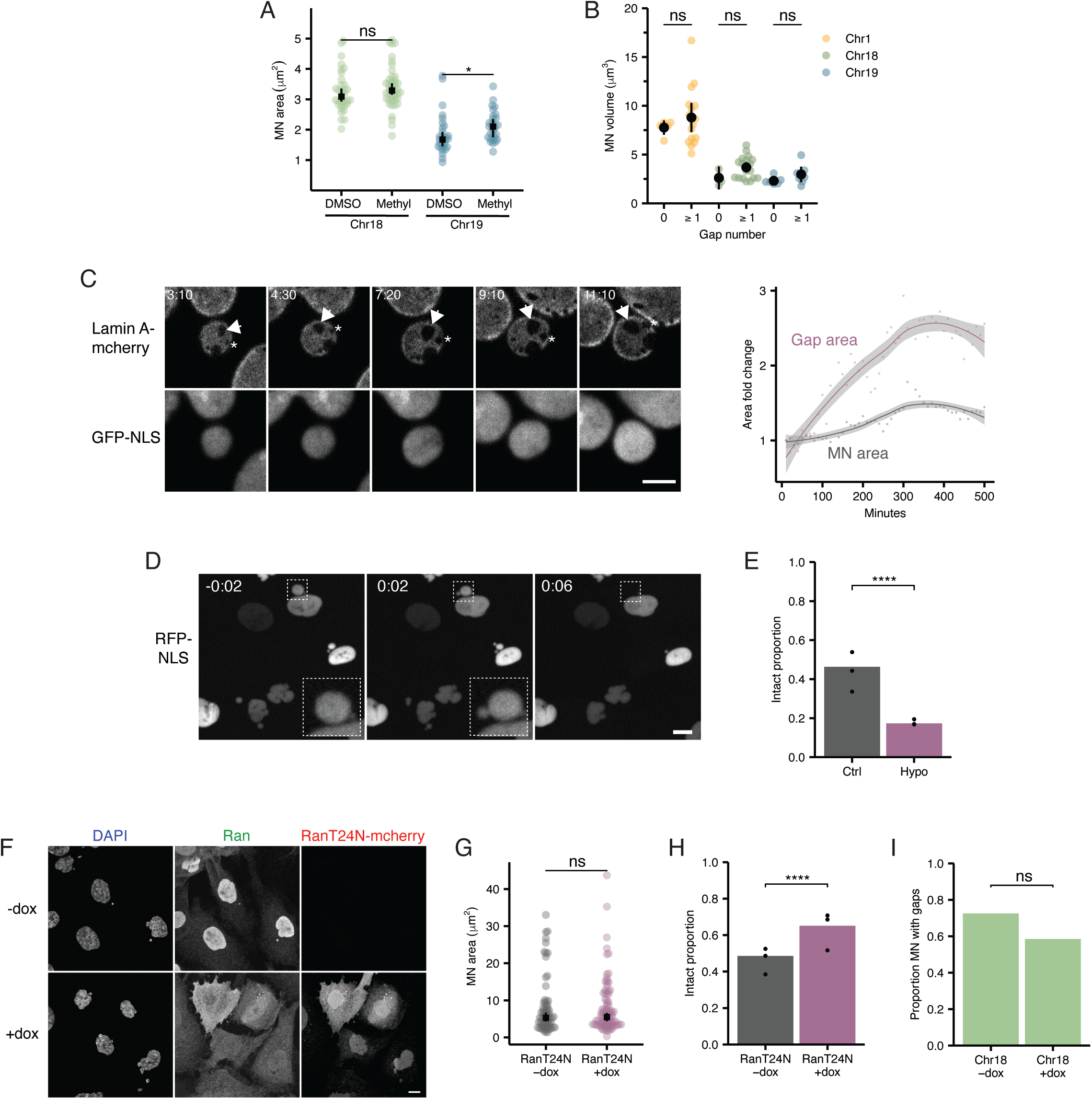
MN growth is required for nuclear lamina gaps and MN rupture. **A.** Maximum projected area of intact MN 24hrs after BAY for single chromosome 18 and 19 MN in RPE1 cells. Chr 19 MN increase in size after methylstat treatment compared to DMSO. Wilcoxon ran sum test, N=3, n=(34, 46, 29, 29). **B.** Comparison of the volume of MN with and without lamina gaps for chr 1, chr 18, and chr 19 MN. Wilcoxon rank sum test, N=3, n=(23, 23, 14) **C.** Example of live imaging from U2OS cells expressing Lamin A-mcherry and GFP-NLS imaged every 10 minutes. Single sections shown. Scale bar = 1um. Lamina gap indicated with an arrowhead was measured throughout imaging while gap indicated by asterisk is not quantified but increases in size. Area fold change for MN area and the arrowhead gap are shown. **D.** Stills from live imaging of RPE1 cells expressing 2xRFP-NLS and treated with hypotonic media diluted 1:2 in sterile water. Cells were imaged every 2 minutes and timepoints from 2 min before hypotonic treatment, and 2 and 6 minutes after are shown. Inset shows MN undergoing nuclear blebbing and rupture. Scale bar = 10um. **E.** MN stability for bulk MN in RPE1 control and cells and after hypotonic swelling for 1 hour. Barnards test, N= 3, n=(525, 433). **F.** Maximum projection images of RPE1 TetOn TRE-RanT24N-mcherry cells treated with 0ng/ul and 100ng/ul dox stained for Ran and DAPI showing nuclear expression of mcherry and loss of nuclear Ran with dox addition. Scale bar = 10um. **G.** Max projected area of intact MN, based on H3K27ac staining, in RPE1 TetOn TRE-RanT24N-mcherry cells treated with 0ng/ul or 100ng/ul dox for 16hrs. Wilcoxon rank sum test, N=3, n=(72,97). **H.** MN stability in RPE1 TetOn TRE-RanT24N-mcherry cells treated with 0ng/ul or 100ng/ul dox for 16hrs. Barnards test, N=3, n= (373,487). **I.** Quantification of nuclear lamina organization in single chromosome 18 MN 24hr post BAY for RPE1 RanT24N-mcherry cells treated with 0ng/ul or 100ng/ul dox. Barnards test, N=3, n=(40, 41). For all graphs, ns p>0.05, * p<0.05, ** p<0.01, *** p<0.001, **** p<0.0001.

We next directly assessed the effect of MN growth on MN stability by quantifying the frequency of intact MN after hypotonic swelling or inhibition of protein import, which is required for nuclear growth^31,32^. RPE1 cells expressing RFP-NLS were incubated hypotonic medium for 1 hr, which is sufficient to significantly increase nuclear area (Fig. S2D, E). Addition of hypotonic medium caused rapid swelling and rupture of a membrane bleb in almost all MN visualized by live-cell imaging (Fig. 2D), which we confirmed by MN rupture analysis in fixed cells (Fig. 2E). Both chr 18 and 19 MN showed a similar loss of stability after swelling (Fig. S2F), consistent with growth being sufficient for MN rupture, regardless of chromosome content.

To inhibit nuclear growth, we inducibly expressed mCherry fused to RanT24N, a GDP-locked Ran mutant that disrupts the Ran gradient and inhibits active protein transport^33–36^. The Ran gradient is required for mitotic spindle and NE assembly^37^. Therefore, RanT24N was expressed only in interphase cells (Fig. S2D). mCherry-RanT24N accumulated in both MN and nuclei and led to decreased levels of Ran in both compartments (Fig. 2F, Fig. S2G,H), indicating a global collapse of the Ran gradient^38,39^. Expression of RanT24N for 16 hrs caused a significant decrease in nuclear size (Fig. S2I). A similar change was not observed in MN (Fig. 2G), likely due to delays in MN growth (see below, Fig. 3, and discussion). Longer RanT24N expression times led to substantial cell death and mitotic disruption, limiting the duration of analysis. However, RanT24N expression was sufficient to significantly increase bulk MN stability, consistent with growth being required for MN rupture (Fig. 2H). To determine whether increased stability correlated with reduced nuclear lamina gaps, we quantified gap appearance on chr 18 MN to reduce noise from analysis of multiple MN types. We found a strong, although not significant, reduction in the proportion of MN with visible lamina gaps (Fig. 2I), strongly suggesting that MN growth contributes to MN rupture by inducing or exacerbating nuclear lamina gaps.

**Figure 3:**
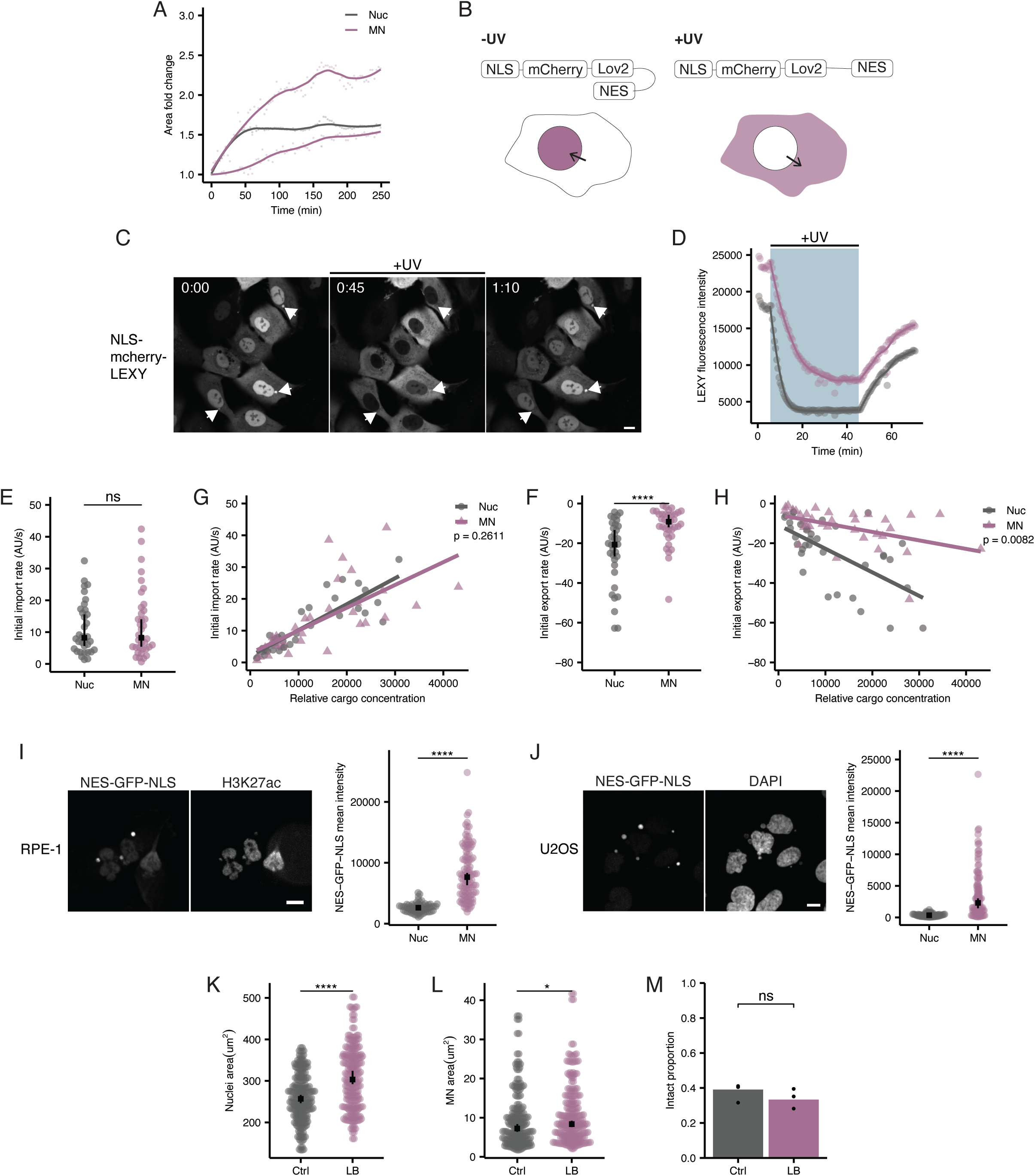
MN have a defect in protein export but not import. **A.** Nuclear area measurements of U2OS cells expressing GFP-NLS synchronized in G2 with 7.5 um R0-3306, released into 100nM BAY, and imaged every 3 minutes. Area of nuclei and MN measured from GFP-NLS channel staring with time 0 at GFP-NLS import after mitosis. **B.** Schematic of NLS-LEXY-mCherry domains. In the absence of UV the NES of LEXY is hidden and the NLS dominates leading to nuclear localization of LEXY. When exposed to UV the NES is uncaged leading to export or LEXY form the nucleus. **C.** Image of single sections from live imaging in RPE1 cells expressing NLS-LEXY-mCherry and H2B-miRFP to identify nuclei and MN during imaging. Cells were images every 30 sec. Stills show cells at the start of imaging without UV, after 45 minutes of UV exposure every 30 sec, and after 25 minutes of imaging in the absence of UV. Scale bar = 10um. **D.** Example for quantification of nuclear and micronuclear mCherry intensity throughout imaging outlined in C**. E.** Nuclear import rates quantified from LEXY import over 25 minutes in the absence of UV in paired nuclei and MN. Paired Wilcoxon rank sum test, N=3, n=34. **F.** Nuclear export rates quantified from LEXY export during 45 minutes of pulsed UV exposure in paired nuclei and MN. Paired Wilcoxon rank sum test, N=3, n=34. **G, H.** Import or export rates compared to the LEXY-mCherry concentration, quantified from cytoplasmic LEXY signal, in RPE1 cells. Analysis of covariance test. **I.** Maximum projection images of RPE1 cells expressing NES-GFP-NLS. Scale bar = 10um. NES-GFP-NLS intensity was quantified for nuclei and intact MN. Wilcoxon rank sum test, N=3, n=(66,112). **J.** Maximum projection images of U2OS cells expressing NES-GFP-NLS. Scale bar = 10um. NES-GFP-NLS intensity was quantified for nuclei and intact MN in U2OS cells. Wilcoxon rank sum test, N=3, n=(73, 117). **K, L.** Maximum projected areas were calculated for nuclei and intact nuclei in RPE1 cells for control and 5hr 20ng/ul leptB treatments. Wilcoxon rank sum test, Nuclei: N= 3, n=(119,125), MN: N= 3, n=(107,126). **M.** MN stability in RPE1 cells for control and 5hr 20ng/ul leptB treatments. Barnards test, N=3, n=(218, 286). For all graphs, ns p>0.05, * p<0.05, ** p<0.01, *** p<0.001, **** p<0.0001.

### MN have a constitutive protein export defect

We next asked what determined growth rate and size in MN. Analysis of MN growth by live-cell imaging found slower rates of MN expansion in early interphase compared to nuclei, consistent with previous observations^19,40^. Despite this, MN frequently exceeded expected sizes, based on area fold change, due to a prolonged period of interphase growth (Fig. 3A), suggesting that nuclear size control is disrupted in MN. Nuclear size is largely determined by protein production in the cytoplasm and nuclear transport rates^41–43^. Since nuclei and MN share a cytoplasm, we focused on potential differences in nuclear transport.

Previous work identified defects in protein import in MN^11,44^. Because this would be expected to inhibit MN growth, we reanalyzed both import and export rates in MN using a small optogenetic reporter, NLS-mCherry-LEXY^45^. This protein contains a weak NLS, an mCherry tag, and a LOV2 domain that binds a strong NES, which can be uncaged by UV light (Fig. 3B). In the absence of UV, NLS-mCherry-LEXY localizes to the nucleus. During UV exposure, the NES is released and the protein is exported. Removing UV recages the NES allowing analysis of import rates in the same cells. RPE1 cells co-expressing NLS-mCherry-LEXY and H2B-miRFP were synchronized, induced to form MN, and imaged every 30s prior to, during, and after pulsed UV exposure (Fig. 3C, D). Imaging durations were determined by defining the average time for mCherry fluorescence in the nuclei and MN to just reach a plateau during UV exposure to limit the effects of prolonged imaging on cell viability. To minimize the effect of the volume differences between MN and nuclei on transport rate analysis, we compared the initial import and export rates^46^ and only between compartments in the same cell.

Surprisingly, we observed no difference in NLS-mCherry-LEXY import rates between MN and nuclei in the same cell (Fig. 3E). To ensure that differences in initial NLS-mCherry-LEXY concentrations in MN or between cells were not affecting these results, we calculated the relative effective transport rate for both nuclei and MN^46^ and similarly observed no difference after adjusting for cargo concentration (Fig. 3G). Instead, we observed a substantial defect in NLS-mCherry-LEXY MN export, with the effective export rate about 2x slower for MN despite a frequently higher starting concentration and a smaller volume that would be expected to reach steady state faster (Fig. 3F). Decreased export rates were observed across all sizes of MN (Fig. S3A), indicating that this is a broadly acting mechanism unlinked to NPC density. In addition, we observed a linear relationship between cargo concentration and transport rate for both MN and nuclei across all concentrations, indicating that NLS-mCherry-LEXY expression is not saturating the transport machinery and therefore unlikely to be affecting these results^46^ (Fig. 3G, H).

We also assessed the relative MN and nuclear NLS-mCherry-LEXY fluorescence intensities at steady state as an additional measure of relative facilitated and passive transport rates. Prior to UV exposure, the NLS-mCherry-LEXY nuclear:cytoplasm (N:C) fluorescence ratio reflects the relative rate of import versus passive efflux. During UV exposure, the N:C ratio reflects the relative active import:export rates^47^. Because their import rates are nearly identical and they share a cytoplasm, we used the MN:nucleus NLS-mCherry-LEXY ratio to analyze changes in the passive export (ratio pre-UV) and facilitated export (ratio during-UV) rates between these two compartments. As expected for a defect in MN export, NLS-mCherry-LEXY MN:nucleus ratios were well above 1 at the end of UV exposure (Fig S3B). MN:nucleus ratios were also slightly above 1 prior to UV exposure (Fig S3B), suggesting that MN have reduced rates of passive diffusion as well. However, we cannot rule out altered reporter binding kinetics in MN versus nuclei as a cause of this difference. To validate our export results, we analyzed the relative nuclear fluorescence of NES-GFP-NLS, which has a strong import and export sequence, in nuclei and MN. Similar to NLS-mCherry-LEXY at the end of UV exposure, NES-GFP-NLS fluorescence was much higher in MN compared to nuclei (Fig. 3I). The difference in NES-GFP-NLS accumulation also increased over time, consistent with decreased protein export rather than decreased DNA decompaction^40^ causing MN accumulation of NES-GFP-NLS fluorescence (Fig S3C). This result was not limited to RPE1 cells, as a similar enrichment of NES-GFP-NLS was observed in U2OS cell MN (Fig. 3J) independent of MN area (Fig. S3D, E).

To assess the severity of this export defect, we asked whether adding leptomycin b (leptB) to inhibit CRM1-dependent export had additive effects on MN size, NES-GFP-NLS accumulation, or MN stability (Fig. S3F). Addition of leptB to interphase cells for 5 hrs was sufficient to increase NES-GFP-NLS nuclear fluorescence (Fig. S3G) and nuclear size (Fig. 3K). We observed a significant, but much smaller effect on MN protein export and size (Fig. 3L, S3G), indicating that CRM1 export is not completely lost, but significantly reduced in MN. Analysis of MN rupture frequency found a small, but not significant decrease in intact MN (Fig. 3M), consistent with a connection between decreased protein export and decreased MN stability. Overall, our data strongly suggest that MN are broadly capable of importing small cargo but have a substantial defect in nuclear protein export, regardless of MN content or size, that enables MN rupture.

### Reduced RCC1 drives nuclear export defects and membrane rupture in MN

Nuclear export rates are determined primarily by exportin/cargo concentrations and Ran gradient strength, with additional regulation by the NPC basket proteins Nup153 and TPR^43,47–51^. To determine the cause of MN export defects, we analyzed the relative recruitment of these factors to MN and nuclei by IF. Because NPC density is directly correlated with MN size^12^ and both basket Nups and exportins are highly concentrated at NPCs^26^, we assessed whether there was additional depletion of Nup153, TPR, and the exportin CRM1 in MN by foci density between these proteins and Nup133, an NPC marker. Similar to Nup133, foci density for all three proteins correlated with MN size, with TPR and Nup153 having the strongest correlation (Fig. S4A-B). Analysis of foci density and intensity, normalized to nuclear levels, found that all three proteins were present at similar levels to Nup133 in MN, with only TPR showing a slight but significant decrease (Fig. 4A-B). These data suggest that reduced recruitment of basket Nups or the major protein exportin CRM1 does not underlie MN export defects.

**Figure 4:**
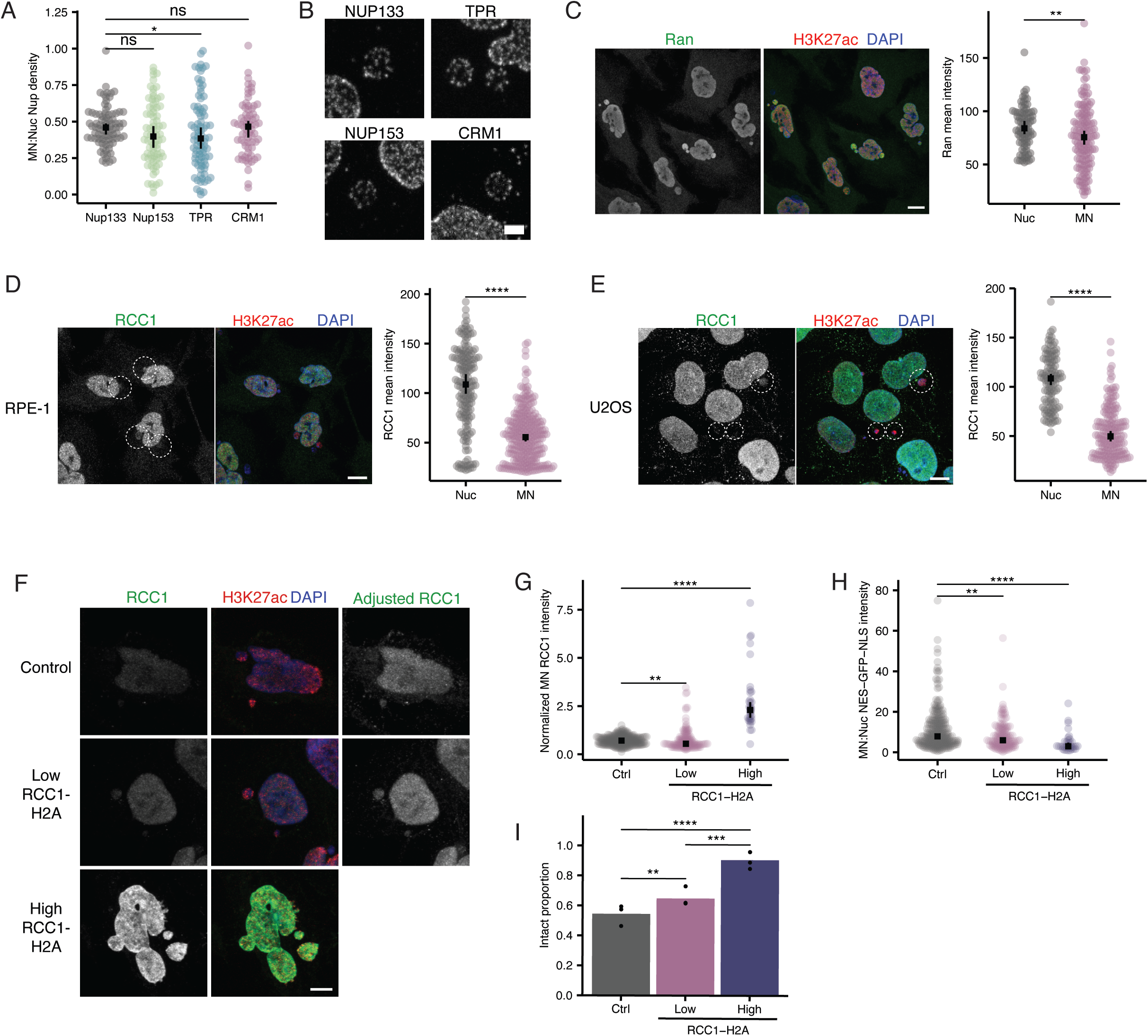
Reduced RCC1 drives nuclear export defects and rupture in MN. **A.** Quantification of NUP density in intact RPE1 cell MN. Wilcoxon rank sum test, N=(3), n=(84,76,84,69). One sample Wilcoxon rank sum comparing MN:Nuc to 1, Nup133 p<0.0001, Nup153 p<0.0001, TPR p<0.0001, CRM1 p<0.0001. **B.** Maximum projections of the top surface of MN in RPE1 cells stained for Nup133, Nup153, TPR, and CRM1. Scale bar =1um. **C.** Maximum projections of Ran in RPE1 cells 24hrs post BAY addition. Quantification of Ran intensity for nuclei and intact MN. Wilcoxon rank sum test, N=3, n=(88, 140). **D.** Maximum projections of RCC1 in RPE1 cells 24hrs post BAY addition. Intact MN indicated with white circles. Quantification of RCC1 intensity for nuclei and intact MN. Wilcoxon rank sum test, N=3, n= (168, 268). **E.** Maximum projected images of RCC1 staining in U2OS cells. Intact MN indicated by white circles. Scale bar= 10um. Quantification of RCC1 intensity for nuclei and intact MN. Wilcoxon rank sum test, N=3, n=(111, 176). **F.** Maximum projections for RCC1 staining in control and RCC1-H2A expressing RPE1 cells, with examples of low and high expressing cells, 24hrs post BAY. Control and low RCC1-H2A cells are also shown with increased contrast so RCC1 is visible in MN. Scale bar= 5um. **G.** Quantification of RCC1 intensity in MN, normalized to median RCC1 intensity for each replicate, in control and RCC1-H2A expressing RPE1 cells. RCC1-H2A cells were categorized as high or low based on nuclear RCC1 levels. **H.** MN:Nuc ratio of NES-GFP-NLS intensity in MN for control and RCC1-H2A expressing cells. **G,H:** Wilcoxon rank sum test, N=4, n=(270, 149, 36). **I.** MN stability in RPE1 cells for control and RCC1-H2A expression. Barnards test. N=3, n=(294, 349, 73). For all graphs, ns p>0.05, * p<0.05, ** p<0.01, *** p<0.001, **** p<0.0001.

In contrast, Ran and RCC1 were significantly depleted in MN compared to nuclei (Fig. 4C-D) in a size-independent manner (Fig. S4C-D). These depletions were also not limited to RPE1 cells as U2OS MN showed a similar strong and broad reduction in RCC1 levels (Fig. 4E, Fig. S4E). To determine whether RCC1 depletion causes MN rupture, we overexpressed RCC1 fused to H2A to drive chromatin enrichment and protein activiation^52^. RCC1 is inhibited by RanT24N, but as this prevents nuclear import as well as export (see Fig. 2F-I), specific effects on export could not be determined. RPE1 cells were nucleofected with RCC1-H2A during release from Cdk4/6i to limit protein expression prior to mitosis (Fig. S4F). RCC1-H2A expression increased RCC1 levels in both MN and nuclei (Fig. 4F), but expression was highly variable between cells due to incomplete nucleofection efficiency. To identify the cell population most affected by RCC1 overexpression, we stratified cells into two groups based on nuclear RCC1 intensity. High expressing cells were defined as those with a mean RCC1 nuclear intensity greater than the highest single cell mean per experiment for control cells. The low expression group is likely a mix of non-nucleofected and low expressing cells.

RCC1 intensity increased in MN in both expression groups compared to controls (Fig. 4G), but only a partial rescue of the relative recruitment of RCC1 to MN versus nuclei was observed (Fig. S4G). Consistent with persistent differences in RCC1 recruitment between MN and nuclei, analysis of relative nuclear export rates by NES-GFP-NLS MN:nucleus intensity ratio quantification found that both expression levels were sufficient to reduce, but not eliminate, the difference in relative export rates between MN and nuclei (Fig. 4H). This reduction was not sufficient to significantly decrease MN size (Fig. S4H, see discussion), but led to a significant increase in MN stability, with high RCC1 expression almost fully rescuing MN rupture (Fig. 4I). Together, these data strongly suggest that RCC1 depletion is a key cause of the defective nuclear export in MN that primes MN for rupture.

### High euchromatin increases MN stability by additional RCC1 loss and decreased import

Our findings suggest that an intermediate reduction of RCC1 in MN enables rupture by causing protein export defects, overgrowth, and nuclear lamina gap enlargement. In contrast, broad inhibition of RCC1 activity by RanT24N expression leads to reduced lamina gaps and increased MN stability, likely by blocking the import pathways required for growth. Recent studies found that RCC1 is more highly recruited to and more active on heterochromatin^53^, leading us to hypothesize that chr 19 MN are more stable than chr 18 MN due to additional reductions in RCC1 levels leading to protein import defects. Consistent with this hypothesis, RCC1 and Ran levels are both reduced in chr 19 MN compared to chr 18 MN (Fig. 5A and S5A). In addition, increasing histone methylation by methylstat led to an increase in RCC1 recruitment to chr 19 MN (Fig. 5B) as well as bulk MN (Fig. S5B). Conversely, increasing histone acetylation on chr 18 MN led to a decrease of RCC1 (Fig. 5B), suggesting that chromatin state is linked to RCC1 recruitment in MN as well as nuclei.

**Figure 5:**
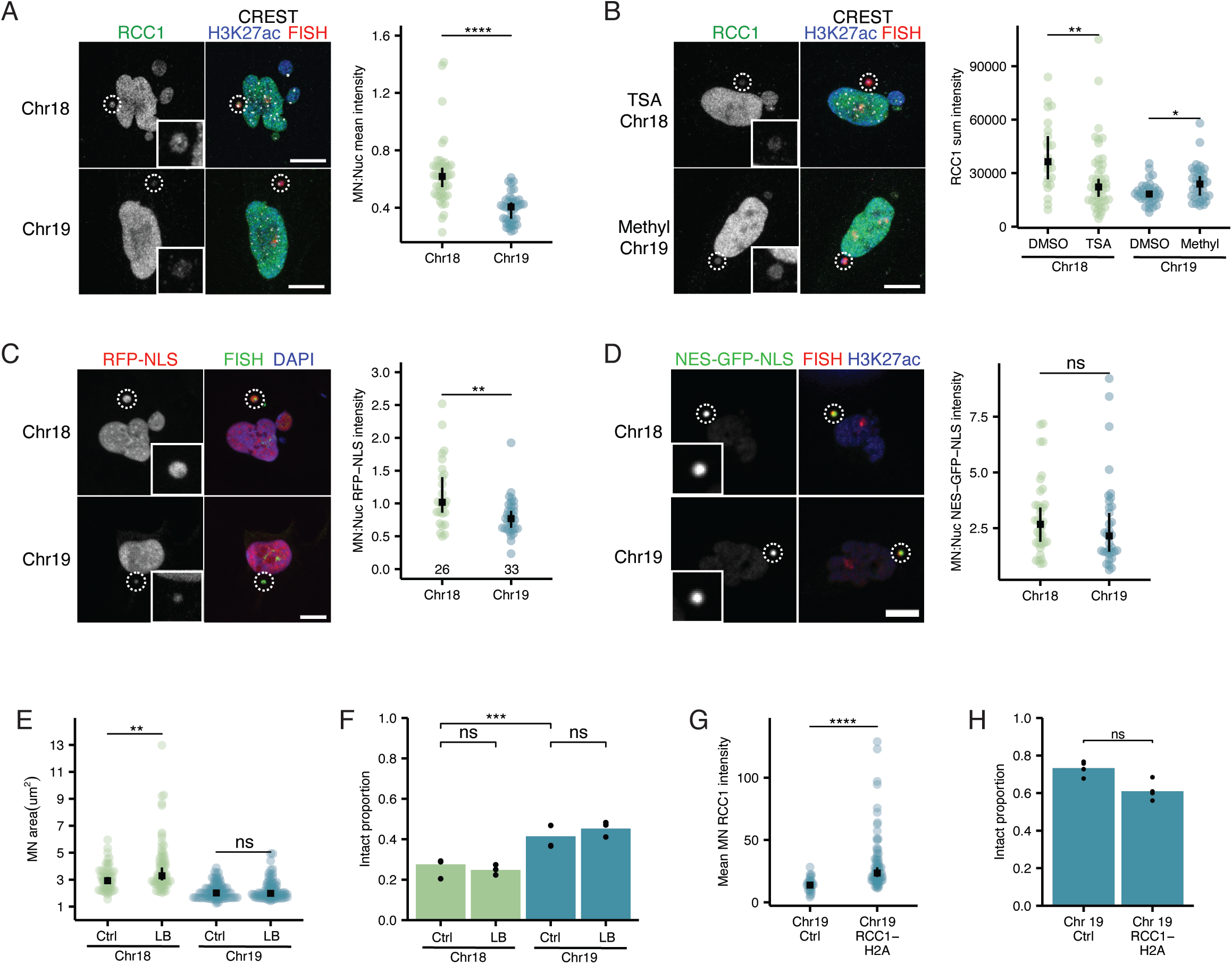
High euchromatin protects chr 19 MN from transport related instability. **A.** Maximum projections of RCC1 in intact single chromosome 18 and 19 MN in RPE1 cells 24hrs post BAY addition. Wilcoxon rank sum test, N= 3, n=(46, 41). **B.** Maximum projections of RCC1 intact single chromosome 18 and 19 MN in RPE1 cells treated with DMSO, 5uM methylstat, or 100nM TSA. Wilcoxon rank sum test, N=3, n=(23, 46, 35, 36). **C.** Maximum projections of intact single chromosome 18 and 19 MN in RPE1 cells expressing RFP-NLS. RFP-NLS intensity was quantified from single sections for nuclei and intact MN. Wilcoxon rank sum test, N=3, n=(26, 33). **D.** Maximum projections of intact single chromosome 18 and 19 MN in RPE1 cells expressing NES-GFP-NLS. NES-GFP-NLS intensity was quantified from single sections for nuclei and intact MN. Wilcoxon rank sum test, N=4, n=(36, 34). **E.** Maximum projected areas for intact single chromosome 18 and 19 MN in RPE1 cells for control and 5hr 20ng/ul leptb (LB) treatments. Wilcoxon rank sum test, N=3, n=(62, 78, 94, 92). **F.** MN stability for single chromosome 18 and 19 MN 24hr post BAY release with 5hr control and 20ng/ul leptB treatments. Barnards test, N=3, n=(261,323,304,243). **G.** Quantification of total RCC1 intensity in single chromosome 19 MN 24hr post BAY for control and RCC1-H2A expressing RPE1 cells. Wilcoxon rank sum test, N=3,4, n=(29,76). **H.** MN stability for single chromosome 19 MN 24hr post BAY in control and RCC1-H2A expressing RPE1 cells. Barnards test, N=3, n=(116, 82). For all graphs, ns p>0.05, * p<0.05, ** p<0.01, *** p<0.001, **** p<0.0001.

To determine whether this additional loss of RCC1 on chr 19 MN was sufficient to disrupt protein import, we compared RFP-NLS MN:nucleus intensity ratios for chr 19 and 18 MN by IF-DNA FISH. We observed a significant decrease in RFP-NLS intensity in chr 19 MN (Fig. 5C), consistent with reduced nuclear import. The other possibility, an increase in passive efflux from chr 19 MN, is highly unlikely due to similarly low NPC densities and Nup recruitments to both types of MN^12,54^. Chr 19 MN, similar to chr 18 MN, accumulated more NES-GFP-NLS than nuclei (Fig. 5D), suggesting that reduced import rates in chr 19 MN were not sufficient to fully suppress the export defect. To directly assess this potential suppression, we asked whether inhibition of protein export by lept B had an additive effect on either size or rupture of chr 19 MN. We first assessed the effect of lept B on chr 18 MN, which has efficient protein import and should be sensitive to export inhibition, and observed a significant increase in chr 18 MN size and a slight, though not significant reduction, in intact MN (Fig. 5E, F), similar to bulk MN (Fig. 3L, M). In contrast, chr 19 MN showed no size increase and no trend towards decreased integrity after leptB incubation (Fig. 5E, F). This is not due to an inability of chr 19 MN to expand, as both chr 18 and 19 MN ruptured at similar frequencies during hypotonic swelling (Fig. S2F). Together these data are consistent with further loss of RCC1 from chr 19 MN inhibiting protein import and suppressing the effects of RCC1-driven defects in protein export.

To directly test whether RCC1 reductions are required for chr 19 MN stability, we analyzed chr 19 MN integrity after overexpressing RCC1-H2A by transient nucleofection. Analysis of RCC1-H2A expression in bulk MN showed that protein export differences were reduced, but not eliminated, between MN and nuclei (Fig. 4H). Therefore, we expected that chr 19 MN would be less stable after RCC1-H2A recruitment. RCC1-H2A expression was sufficient to increase RCC1 levels in chr 19 MN (Fig. 5G) and led to a consistent, albeit not significant, decrease in chr 19 MN stability (Fig. 5H). Thus, our results suggest that increasing RCC1 levels in most MN increases their stability by rescuing nuclear export, but that increasing RCC1 levels on small euchromatic MN has the opposite effect: enhancing the export defect and decreasing stability by improving protein import.

### MN rupture timing correlates with DNA damage signatures

Our data define an order of MN rupture based on chromatin content leading to differences in RCC1 and B-type lamin recruitment. Previous work suggested that DNA damage patterns on micronucleated chromatin can be linked to rupture timing, with APOBEC-dependent hypermutation occurring throughout the cell cycle and more extensive double stranded DNA breaks occurring only after DNA replication initiation^5,6,55–57^. Based on this model, we hypothesized that rupture timing could be a key determinant of DNA damage signatures from MN rupture events. To test this hypothesis, we analyzed data collected from the Pan-Cancer Analysis of Whole Genomes (PCAWG) Consortium, which contains whole genome sequencing data of chromothripsis events from 2,658 tumors across 38 cancer types^58^. We divided chromosomes into 3 categories: early rupturing (chr 13, 18, and 21), mid rupturing (chr 7-12, 14-16, 20, and 22), and late rupturing (chr 1-6, 17, 19, X) based on our previous data (Fig. 6A)^12^. We then filtered for chromosomes that were annotated as “high confidence chromothripsis” or “no chromothripsis” by ShatterSeek^59^. Since chromothripsis can arise from complex ongoing, including chromatin bridges^60^, we removed chromothripsis events annotated as occurring “with other complex events” to enrich our dataset for those likely associated with MN. This yielded a dataset of 37 early rupture, 128 mid rupture, and 166 late rupture chromothripsis events across all available cancer types (Table S1, S2). The n values for each rupture timing class likely reflects the number of chromosomes included in each category rather than biases in chromothripsis annotations as all annotations showed a similar proportion of each rupture timing category (Fig. S6A, B).

**Figure 6.**
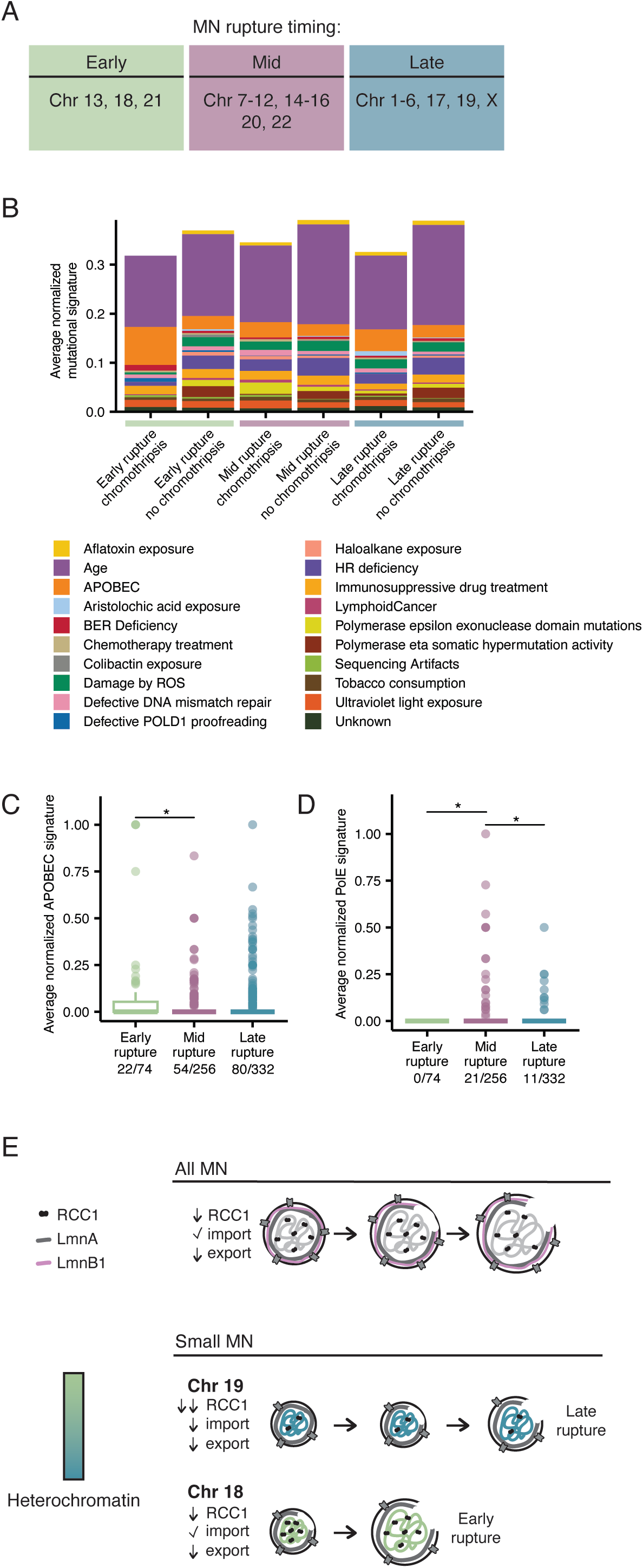
APOBEC and POLE signatures in chromothripsis events are correlated with MN rupture timing. **A.** MN rupture timing classification based on chromosome length, LAD density, and previously analyzed rupture frequencies in RPE1 cells. **B.** Mutational signatures are broadly present and identifiable in all rupturing timing chromosome classes, in the presence (“chromothripsis”) and absence (“no”) of chromothripsis events. Chi-square test, p=ns. **C,D.** Survey of mutational signatures identifies APOBEC and the POLE mutation signatures as enriched in high confidence non-complex chromothripsis events in early and mid rupture chromosomes, respectively. Number of non-zero datapoints out of the total are indicated under each group. Each mutation signature includes 2 unique signatures; APOBEC signature (SBS 2 and SBS 13) and PolE signature (SBS10a and b). (Anova, APOBEC p=0.0166, PolE p=0.005. Tukey honest significant difference test used for pairwise differences. n= 74, 256, 332. **E.** Model of MN rupture due to transport induce growth in all MN and small MN. For all graphs, ns p>0.05, * p<0.05, ** p<0.01, *** p<0.001, **** p<0.0001.

We next downloaded all single nucleotide variant (SNV) calls from the PCAWG Consortium^61^ and used Sigprofiler^62^ to assign mutational signatures to each one. Mutational signatures were broadly present in all three rupture timing categories regardless of chromothripsis status, indicating that differences in chromosome characteristics are not a major contributor to mutation type (Fig. 6B). Further analysis of mutational signatures by rupture timing category and chromothripsis status identified a significant enrichment of APOBEC signatures in early rupture chromosomes with chromothripsis (ANCOVA, p=0.0166, Fig. 6C, Table 1). Importantly, this enrichment was not observed in “no chromothripsis” sequences (Fig. S6C). These results are in line with proposed mechanisms of MN DNA damage during G1 phase^63,64^ and with a general enrichment of APOBEC signatures in tumors with chromothripsis^59,65^ and suggest that rupture early in the cell cycle can bias MN chromatin towards hypermutation. Surprisingly, we also identified an enrichment of signatures associated with DNA polymerase POLE hypermutation on mid-rupture chromosomes (ANCOVA, p=0.005, Fig. 6D, Table 2) that, similar to APOBEC, was not observed in non-chromothripsis chromosomes (Fig. S6D). None of the patient samples are annotated as having a POLE SNV associated with hypermutation^66^, suggesting that these signatures may be arising from a mechanism related to nuclear function disruption in intact MN^67^.

**Table 1.** APOBEC mutational signature data. For figure 6, APOBEC signature values with chromosome rupture timing and sample IDs from the PCAWG dataset.

**Table 2.** POLE mutational signature data. For figure 6, POLE signature values with chromosome rupture timing and sample IDs from the PCAWG dataset.

## Discussion

MN formation and rupture are considered a major driver of large and rapid genome and signaling changes that promote tumorigenesis^68^. A hallmark of MN are large gaps in the nuclear lamina that become the sites of membrane rupture^69,70^. Similar gaps in the nuclear lamina in the nucleus arise in response to disease-associated changes in NE protein expression and chromatin structure^71^. However, the mechanisms underlying nuclear lamina gap initiation and expansion remain unclear. In this study we demonstrate that MN have a constitutive defect in protein export due to reduced levels of the transport factor RCC1 and that this export defect is a key driver of nuclear lamina gaps and membrane rupture (Fig. 6E, top). However, this pathway can be modified by the chromatin content of the MN. Previously, we found that large MN containing at least one long human chromosome or multiple short ones efficiently recruited lamin B1^12^, likely due to the larger areas of flat membrane ^72,73^, which inhibits lamina gap formation^69,74^. Here we demonstrate that low levels of histone methylation in small MN also inhibit nuclear lamina gap formation, but due to additional defects in RCC1 recruitment that result in reduced micronuclear import and membrane growth (Fig. 6E, bottom). Together these findings define a broad mechanism for MN rupture that explains the prevalence of nuclear lamina disorganization in these compartments and known differences in MN rupture frequencies and timing across systems.

### Mechanisms of MN nuclear lamina gap formation and membrane rupture

Our data demonstrate that nuclear membrane expansion is required for the appearance of nuclear lamina gaps, but we do not know whether expansion is sufficient to initiate them. Other models for lamina gap initiation are improper resolution of microtubule-chromatin contacts during anaphase^75^ and strain on NPC/nuclear lamina interactions^76^. Both could contribute to the frequency of nuclear lamina gaps in MN and additional modeling of the nuclear lamina network and mechanics in MN could yield new insight into the formation of these defects across nuclear compartments.

Our results also suggest that nuclear lamina assembly is highly regulated and decoupled from membrane growth in MN. This is consistent with a longstanding model where interphase assembly of “non-core” NE proteins, including the lamin B meshwork and NPCs, is regulated by a different set of kinases than nuclear growth^77^. Nuclear lamina assembly is also inhibited or severely delayed on membrane blebs that form in the nucleus^78–82^, suggesting that this decoupling is not MN specific. Interestingly, nuclear membrane blebs fail to recruit B-type lamins despite the presence of active kinases proposed to drive rapid NPC and nuclear lamina assembly in early interphase^77,83^. Thus it is unclear whether large nuclear lamina gaps in MN represent an unappreciated aspect of normal interphase nuclear lamina assembly or a lack of nuclear lamina assembly factors from MN.

Analyses of both MN and nuclear membrane rupture demonstrated that nuclear lamina gaps are often required, but not sufficient, for membrane rupture^69,84–88^. Similarly, we observed extensive expansion of MN nuclear lamina gaps without rupture, suggesting that they enable membrane disruption but cannot trigger it. What then is the rupture trigger? In the nucleus, rupture is often driven by mechanical force, indicated by the formation of a membrane bleb prior to loss of compartmentalization^71^. We observe membrane blebs in MN frequently during hypotonic swelling (Fig. 2D), but only rarely during normal rupture, suggesting that spontaneous MN rupture is seldom triggered by mechanical force^16,80^. Recent work has found that activation of ATR^24^ or Chmp7^89–91^ in MN can trigger rupture in different subsets of MN. Chmp7 accumulation has been linked to export defects in nuclei and MN^91–93^, and determining how RCC1 activity impacts Chmp7 and ATR in addition to its effects on nuclear lamina organization will be critical to defining the full mechanism of MN disruption.

LADs have been proposed to be essential for proper chromatin organization, gene expression, and mechanoprotection^94,95^. Surprisingly, we do not observe broad differences in peripheral H3K9me2 enrichment between chr 18 and chr 19 MN, despite an almost 2.5-fold enrichment of LAD density on chr 18 in RPE1 nuclei^12,96^. This could be related to the overall enrichment of H3K9me2 on micronucleated chromatin^19,20^, but our analysis of chromosome-specific histone modifications finds that many differences observed in nuclei are preserved in MN, consistent with many classes of histone modifiers localizing to MN^97,98^. Instead, our results suggest that, despite reduced levels of B-type lamins and NPCs in small MN, the available nuclear lamina is sufficient to properly organize heterochromatin at the periphery. Whether this tethering contributes to the mechanics or rupture timing of small MN remains to be determined.

Nuclear volume is strongly correlated with cytoplasm volume, *i.e.* the N:C ratio, with growth rates largely determined by the rate of nuclear import^99^. In mammalian cells, nuclear expansion is limited by the nuclear lamina^100^ and ER availability^42,101^, with a minimum nuclear volume set by forces generated by the chromatin^32,41,102^. We propose that the slow growth rate of MN leads to a strong correlation between MN size and chromatin amount^12^, especially during G1. This limits the capacity for MN to shrink, which appears in our data as non-significant MN size reductions in conditions expected to inhibit growth, and may underlie observations of DNA overcompaction in MN^40,97^.

Facilitated nuclear transport rates are largely set by cargo concentration and Ran gradient strength^47^. Previous analysis of micronuclear import rates led to contradictory models of MN transport that ranged from a complete inability to import small proteins to reduced transport only of large cargoes^16,67,69^. Our analysis of NLS-mCherry-LEXY transport demonstrated that MN containing whole chromosomes do not have an import defect, but instead fail to properly export even small cargo. The exception being highly euchromatic small MN, which we find have an import defect, and potentially very small MN formed around chromatin fragments, which were excluded from this study due to their extremely rapid rupture in our experiments (data not shown). The lower concentration of RCC1 and Ran we observed in bulk MN are consistent with both a broad export defect and slower import of large cargo, as both processes are highly sensitive to lower levels of RanGTP^51,103,104^. Thus, we anticipate that rescuing RCC1 levels in MN will also improve import of large proteins and complexes, which may independently contribute to the increased membrane and nuclear lamina integrity we observed through rescue of other nuclear processes.

Competition with the nucleus for NE proteins has been proposed as a key driver of post-mitotic protein recruitment defects in MN^16^, but our results suggest that competition for transport cargo may be a more critical contributor to MN rupture. First, nuclei may compete with MN for RCC1. We find that overexpressing histone associated RCC1 is not sufficient to fully erase the differences in RCC1 recruitment between MN and nuclei (Fig. 5), suggesting that additional mechanisms, likely acting in both mitosis and interphase^105,106^, regulate its recruitment. In contrast, nuclei competition for cargo could delay nuclear lamina disruption by reducing MN import. Whether this competition prevents accumulation of specific protective proteins into the MN, such as nuclear lamina assembly factors, remains to be determined. Interestingly, a recent report suggests that MN can accumulate the essential import protein importin α^107^, consistent with an RCC1-dependent export defect^104^. This suggests that MN formation could affect transport efficiency in the main nucleus, and that MN rupture may in fact be beneficial for the cell.

### The influence of MN rupture timing in cancer development

MN rupture is strongly linked to increased aneuploidy, cancer genome rearrangements, and invasive and inflammatory signaling^68^. Our work suggests that the timing of MN rupture with regards to the cell cycle may also determine likelihood and type of hypermutation. We find that chromothripsis-associated APOBEC signatures are specifically enriched on chromosomes that induce early MN rupture and that POLE-like hypermutation is depleted from this group and unexpectedly enriched on MN that are more likely rupture during DNA replication. Currently the contribution of whole chromosome MN rupture to cancer genome evolution is unclear. However, our findings have significant implications for two major types of missegregation identified from cancer genome sequencing – extrachromosomal DNAs (ecDNAs) and chromosome arm fragments^108,109^. ecDNAs are circular pieces of DNA that are a major source of oncogene amplification and frequently form large MN, especially after induced DNA damage, capable of transcription and replication^110–114^. Micronucleation of ecDNAs is thought to contribute to their elimination and inducing early rupture of these MN may be effective to further suppress ecDNA replication and inheritance. Chromosome arm alterations are extremely prevalent in cancer genomes and induced arm fragments are frequently missegregated into MN^109,115^. We found MN containing acentric fragments to be highly unstable in our assays, suggesting that they rupture early, similar to other small MN. Whether this also increases the probability of APOBEC hypermutation or leads to enhanced cGAS and invasive signaling remains to be determined. Overall, our findings suggest that MN rupture timing is an important factor determining the pathways of cancer genome evolution across numerous tumor types.

## Materials and Methods

### Cell lines and culture methods

hTERT-RPE-1 (RPE1) cells were grown in DMEM/F12 (Gibco) supplemented with 10% fetal bovine serum (FBS, vol/vol, Gibco), 1% penicillin-streptomycin (pen-strep, vol/vol, Gibco), and 0.01 mg/ml hygromycin (Sigma-Aldrich) at 5% CO_2_ and 37°C. U2OS cells were cultured in DMEM (Gibco) supplemented with 10% FBS and 1% pen-strep at 10% CO_2_ and 37°C. RPE1 cells expressing TetON were cultured in tet-free FBS (GeminiBio). RPE1 NES-GFP-NLS cells were made by transduction of NES-GFP-NLS lentivirus, followed by selection with blasticidin (Invivogen) and FACS enrichment for GFP positive cells. RPE1 H2B-emiRFP cells were made by transduction of H2B-emiRFP lentivirus followed by selection with G418 (Thermo-Fisher). RPE1 2xRFP-NLS were made by transduction of 2xRFP-NLS lentivirus, followed by selection with blasticidin and FACS enrichment for high expressing RFP positive cells. RPE-1 TetON cells were made by transfection of pLVX-Neo:Eto.1 (TetOn) lentivirus and selection with G418. RPE1 TetOn TRE-RanT24N-mcherry cells were then made by lentiviral transfection of pLVXE-Blast:TRE-RanT24N-mcherry into RPE1 TetOn cells and selection with blasticidin. U2OS GFP-NLS Lamin A-mCherry cells were previously described^116^. U2OS NES-GFP-NLS cells were made by transduction of pLVXE-Blast:NES-GFP-NLS lentivirus, followed by selection with blasticidin.

MN were induced in synchronized RPE1 and U2OS cells by arresting cells in G1 by addition of 1 μM PD-0332991 isethionate (Cdk4/6i; Sigma-Aldrich) for 24 h, washed 3 times with 1x PBS, released into fresh medium containing 100 nM BAY-1217389 (Mps1i; Thermo Fisher), and fixed 24 h later, unless noted in figure. Methylstat (histone demethylase inhibitor; Abcam), TSA (HDACi; Cell Signaling Technologies), and DRB (RNA Pol II inhibitor; Sigma), were added at 5 µM, 100 nM, and 100 µg/mL, respectively, at 20 h post Mps1i addition for 4 h before fixation. Control cells were incubated in equivalent amounts of DMSO. For lamina gap analyses, cells were fixed at 20 h post Mps1i addition. For MN and lamina gap growth analyses, U2OS GFP-NLS Lamin A-mCherry cells were incubated in 7.5 µM RO-3306 (Cdk1i; Sigma Aldrich) for 16.5 h, washed 7 x with warm media, and released into media containing 100 nM BAY-1217389. Live imaging started less than 1 h after release from synchronization.

Hypotonic swelling of RPE1 cells was performed by incubating cells 22 h post Mps1i addition in Mps1i-containing media diluted 1:2 with sterile water. For live imaging, imaging was started 30 m prior to diluting the media. For IF-DNA FISH, cells were incubated in hypotonic medium for 1 h, then Mps1i-containing isotonic medium for 1 h to restore chromatin structure before fixation.

RanT24N-mCherry was induced in RPE1 tetON cells by adding 100 ng/µl doxycycline (Clontech) 8 h post Mps1i addition for 16 h prior to cell fixation. Leptomycin B (Sigma Aldrich) was added at 20 ng/uL to RPE1 cells 19 h post Mps1i addition and imaged immediately for live-cell imaging or fixed after 5 h for IF-DNA FISH. Control cells for leptomycin B treatment were incubated in an equal volume of 100% ethanol.

### Plasmids

2xRFP-NLS was generated by inserting mCherry-NLS-tagRFP (G-Block, IDT) into pLVXEblast (pLVXE-IRES-puro (Clontech) with puroR replaced by blastR) cut with BamHI - EcoRI by Gibson assembly. The NLS sequence is PPKKKRKV. TRE-RanT24N-mCherry was constructed by PCR and ligation of RanT24N-mCherry from pTK21 into pLVXTight (Clontech) cut with BamHI - MfeI and modified to replace the CMV promoter with Ef1a and puroR with blastR. pTK12 was a gift from Iain Cheeseman (Addgene plasmid # 37396). pLVX-Neo:Eto.1 was a gift from the Fred Gage lab (Salk Institute) and is pLVX-Tet-On Advanced (Clontech/Takara) with the CMV promoter replaced with the EF1a promoter. H2B-miRFP was generated by PCR from pH2B- miRFP703 (a gift from Vladislav Verkhusha, Addgene plasmid # 80001) and ligation into pLVXEneo (pLVXEblast with blastR replaced with neoR) cut with NotI - MluI. NLS-mCherry-LEXY (pDN122) was a gift from Barbara Di Ventura & Roland Eils (Addgene plasmid # 72655). NES-GFP-NLS was constructed by PCR and ligation of NES-GFP-NLS from pcLumioDest:NES-GFP-NLS into pLVXE-Blast cut with ClaI - NotI. pcLumioDest:NES-GFP-NLS was a gift from the Martin Hetzer lab (ISTA, Austria) with the previously described NLS sequence and the NES sequence LQLPPLERLTL. pLVXE:2xGFP-NES was described previously^12^. pLVXE-Blast:RCC1-FLAG-H2A was generated using Vector Builder (vector ID is VB240110-1335jye).

### Plasmid transfection

After incubation in 1 µM Cdk4/6i for 24 h, RCC1-FLAG-H2A was transfected into RPE1 cells using a 4D nucleofector (Lonza) and the SE cell line 4D-Nucleofector X Kit S (Lonza) according to kit directions. 200,000 cells were resuspended in buffer SE with 400 ng plasmid and electroporated using program DS-138. Cells were plated into pen-strep free media containing 100 nM BAY and fixed 24 h later. NLS-mCherry-LEXY was transfected into asynchronous RPE1 H2B-miRFP cells using the same conditions, plated into pen-strep free media, and treated with 100 nM BAY for 16 h before imaging.

### NLS-mCherry-LEXY live cell imaging

RPE1 H2B-miRFP cells nucleofected with NLS-mcherry-LEXY were plated into a 2-well glass bottom ibidi plate in pen-strep free media. After 8 h of recovery, cells were incubated in medium containing 100nM BAY for 16 h in the dark. Images were acquired using a 40×/1.3 Plan Apo objective on an automated Leica DMi8 microscope outfitted with a Yokogawa CSU spinning disk unit, Andor Borealis illumination, an ASI automated stage with Piezo Z-axis, and an Okolab controlled environment chamber (humidified at 37°C with 5% CO2). Images were captured with an Andor iXon Ultra 888 EMCCD camera using MetaMorph software (version 7.10.4; Molecular Devices). Cells on the microscope were kept in the dark for at least 15 m prior to imaging and throughout the imaging period. Images were captured every 30 s for 5 m using first only the 568 and 647 lasers, then for 45 min with the 405, 568, and 647 lasers, and then for 20 min with only the 568 and 647 lasers. The duration and intensity of the pulsed 405 exposure was determined experimentally and extended until the nuclear and MN mCherry levels had just reached plateaued to limit cell death.

LEXY intensity was quantified by manually defining nuclei and MN ROIs on the H2B-miRFP channel images in FIJI for each timepoint and measuring the mean intensity on the LEXY-mCherry channel. Only cells where the nuclei showed the expected changes in LEXY intensity during import and export cycles were analyzed. Only MN that remained intact throughout imaging were analyzed. Cytoplasm LEXY intensity was quantified by manually drawing the largest possible ROI within the cytoplasm defined by LEXY-mCherry that did not include obvious dark organelles and then averaged for the first 5 images of the initial 2-channel imaging (initial) and the last 5 time points of 3-channel imaging (export). Similarly, initial and plateau compartment areas, mean intensities, and intensity ratios were obtained by averaging values from the first 5 frames of 2-channel imaging and the last 5 frames of 3-channel imaging.

Initial import rates were quantified by plotting mean fluorescence intensity versus time for all compartments from the start of the last 2-channel imaging period and fit to a one phase association curve in Prism (version 10.0.3) using the equation I_t_ = I_max_ * (1-e^-kt^), where I = fluorescence intensity, I_max_ = fluorescence intensity at the plateau, and t = time. From this equation, the mean fluorescence value of I_max_ and rate constant (k) were derived. The initial import rate was calculated from the curve fit values using the equation: initial rate = k*I_max_. Import rate values were discarded if the curve fit was poor (R^2^ < 0.9, no convergence on either I_max_ or k values), the curve decreased for a large part of imaging, or the plateau values were visually noisy due to focal drift. In addition, only cells where rate values were obtained for both the nucleus and MN were kept to control for differences in photobleaching rates between cells.

Derivation of initial export rates from a one phase dissociation curve fit (initial rate = -k*I_0_) significantly overestimated the rate for many curves due to noisy predictions of I_0_ and led to large differences in rate estimation between similar curves. Therefore, linear regression analysis was used as described in ^117^. LEXY-mCherry mean intensity versus time was plotted starting with the first 3-channel image and a line fit by linear regression through the first 3 time points. The slope of this line is equivalent to the initial export rate. Some compartments did not initiate export immediately. In these cases, the linear regression was shifted forward by 1 timepoint until a negative slope was obtained. Initial export values were excluded if a linear regression with an R^2^ > 0.9 could not be obtained within 3 time frame shifts or if either the nuclear and MN rates could not be determined in a cell. Statistical differences between MN and nuclear initial transport rates were calculated using the Wilcoxon matched-pairs signed rank test in Prism.

Relative effective import rates were calculated as follows: initial import rates for MN and nuclei were plotted versus the cytoplasmic LEXY-mCherry mean intensity for that cell, which indicated the relative initial cargo concentration between cells. The slope of the line fit by linear regression is equivalent to the effective import rate^117^. These fits show the expected correlation between increased cargo concentration and increased import rate indicating that expression of LEXY-mCherry is non-saturating. Relative effective export rates were calculated by plotting the initial export rates versus the average initial MN and nuclear mean intensity, which indicated the relative initial cargo concentration between compartments in the same cell, and performing a similar linear regression analysis. Statistical differences between the effective import and export rates of MN and nuclei were calculated using an analysis of covariance (ANCOVA) test in Prism.

### Immunofluorescence

RPE1 and U2OS cells were grown on poly-L-lysine coated coverslips and fixed in 4% paraformaldehyde (16% pfa (Electron Microscopy Sciences) diluted in 1x PBS) for 10 m at RT. For RCC1 labeling, cells were pre-extracted for 15 s in 0.1% Triton X-100 (Sigma-Aldrich) in 1xPBS at RT, then fixed in 100% MeOH for 10 m at -20°C. After fixing, coverslips were blocked in PBSBT (3% bovine serum albumin (BSA, Sigma-Aldrich), 0.4% Triton X-100, 0.02% sodium azide (Sigma-Aldrich),1× PBS) for 30 m RT, then in the primary antibody for 30 m RT, then secondary antibody for 30 m RT (except where noted below) with 3 x 5 m RT washes in PBSBT between antibodies. After the final secondary antibody, coverslips were washed twice in PBST, incubated in DAPI (1 µg/ml in PBS; Roche) for 5 m RT, washed once in diH_2_O, and mounted in Vectashield (Vector Labs) or Prolong Gold (Life Technologies).

Primary antibodies used: mouse anti-H3K27ac (1:250; 1 h; 39085, Active Motif) rabbit anti-H3K27Ac (1:500; ab4729; Abcam), rabbit anti-H3K27me3 (1:500; MA511198; Thermo Fisher), rabbit anti-H3K9me2 (1:250; AM39041; Active Motif), rabbit anti-Phospho-Rpb1 CTD (Ser2) (1:500; 13499; Cell Signaling Technology), human anti-CREST (1:100; 1 h; 15-234; Antibodies Incorporated), rabbit anti-LBR (1:500; ab32535; Abcam), rabbit anti-Lamin A (1:500, O/N at 4°C; L1293; Sigma Aldrich), mouse anti-Ran (1:500; O/N at 4°C; MA511198; Thermo Fisher), mouse anti-RCC1 (1:200; 2 h, refix with 4% pfa; sc-376049; Santa Cruz Biotechnology), rabbit anti-Nup133 (1:100; ab155990; Abcam), rabbit anti-Nup153 (1:100; ab96462; Abcam), rabbit anti-TPR (1:500; ab84516; Abcam), rabbit anti-CRM1 (1:1000; 46249; Cell Signaling Technology), RFP-boost (1:1000; 1 h; rb2AF568; Chromotek), rabbit anti-FLAG (1:500; F7425; Millipore Sigma). All primary antibodies were diluted in PBSBT.

Secondary antibodies used: AF647 goat anti-human (1:1000; A-21445; Thermo Fisher), AF488 goat anti–mouse (1:1000; A-11029; Thermo Fisher), AF405 goat anti-mouse (1:500; 1 h; A-31553; Thermo Fisher), AF594 donkey anti-rabbit (1:500; 711-585-152; Jackson ImmunoResearch), AF568 goat anti-mouse (1:1000; A11031; Life Technologies), AF488 goat anti–rabbit (1:1000; A-11034; Thermo Fisher), AF568 goat anti-rabbit (1:1000; A11036; Life Technologies), AF405 goat anti-rabbit (1:500; 1 h; A-31556; Thermo Fisher). All secondary antibodies were diluted in PBST.

### IF-DNA FISH

For IF-DNA FISH experiments analyzing RFP-NLS, NES-GFP-NLS, or Ran, cells were first fixed in 4% pfa for 10 m at RT. For all other IF-DNA FISH experiments, cells were first fixed in 100% MeOH for 10 m at - 20°C. IF was performed as above, stopping before DAPI labeling. Coverslips were re-fixed in 4% PFA for 5 m RT, washed 2 x 5 m with 2x SSC (Sigma-Aldrich), and permeabilized in 0.2 M HCl + 0.7% Triton X-100 for 15 m RT. Coverslips were then washed 2 x 5 m with 2x SSC, denatured in 50% formamide (EMD Millipore), 2x SSC for 1 h, washed 2 x 5 m with 2x SSC, then inverted onto 3–5 µL of Spectrum Orange or Green XCE or XCP probes (MetaSystems) and sealed with rubber cement. Probes and targets were co-denatured at 74°C for 3 m and hybridized 2 h – O/N at 37°C in a humidified chamber (2 h for XCE probes and O/N for XCP probes). Cells initially fixed in MeOH were then washed once in pre-heated 0.4x SSC at 74°C for 5 m then 2 x 5m in 2x SSC + 0.1% Tween-20 (Sigma-Aldrich). Cells initially fixed in PFA were washed once in pre-heated 0.25× SSC buffer at 74°C for 5 m then 2 x 5 m in 4x SSC + 0.1% Tween-20. Coverslips were then incubated in DAPI (1 µg/mL in PBS) and mounted in Vectashield (Vector Labs) (for analysis of MN morphology or protein recruitment) or Prolong Gold (Life Technologies) (for MN rupture analysis).

### Microscopy

Unless noted below, images were acquired with a Leica DMi8 laser scanning confocal microscope using the Leica Application Suite (LAS X) software and a Leica ACS APO 40×/1.15 Oil CS objective or a Leica ACS APO 63×/1.3 Oil CS objective. Z-stacks were acquired with the system optimized step size for each objective except for: MN rupture only analysis (0.5 µm step), nuclear lamina gap analysis in RanT24N expressing cells (0.2 µm step, with Lightning Deconvolution).

Nuclear lamina gap analysis was performed on cells imaged using a Leica TCS Sp8 with STED super-resolution using a 775 nm pulsed laser, LAS X (version 3.5.7.23225), and a Leica HC PL APO 100×/1.4 Oil CS2 objective. Before image acquisition the STED and confocal beams were manually aligned using FluoSpheres mounted in Prolong Gold. Images were acquired at ∼20 nm pixel size for a resolution of ∼50 nm in the xy plane, and a white light laser was tuned to 405 nm (DAPI), 488 nm (H3K27ac), 556 nm (FISH), 594 nm (Nup133 or lamin A), and 647 nm (CREST) wavelengths. Post-acquisition, images were deconvolved using LAS X Lightning.

RCC1 intensity analysis was performed on cells imaged on a Leica Stellaris with a Leica HC Plan Apo 63x /1.40 Oil CS2 objective, and LAS X (version 4.0.2). For all images, post-acquisition image processing was limited to cropping the image and adjusting levels with FIJI or Adobe Photoshop to make use of the entire histogram spectrum. False colors for channels were changed through the arrange channels function in Fiji ^118^.

### Time lapse imaging

Time lapse images were acquired using a 40×/1.3 Plan Apo objective on the Leica DMi8 microscope with Yokogawa CSU spinning disk unit described in the LEXY live cell imaging section. For the hypotonic treatment imaging in RPE1 RFP-NLS cells a single section was imaged at 2 min intervals for 2 hrs. For post-mitotic MN and nuclei growth imaging, U2OS GFP-NLS Lamin A-mCherry cells were imaged with a z-step size of 0.5 µm at 3 m intervals. For leptomycin B treatment imaging in RPE1 NES-GFP-NLS cells, cells were imaged as single slices at 5 m intervals for 8 h for control (used for NES-GFP-NLS intensity measurements in MN over time) and leptomycin B treated cells.

Nuclear lamina gap growth images of U2OS GFP-NLS Lamin A-mCherry cells were acquired using a 100x/1.4 (oil) objective on an Andor Dragonfly 200 High Speed Confocal Microscope outfitted with a humidified environmental chamber. Images were captured with an Andor iXon 888 EMCCD using Fusion software and were taken every 10 m as z-stacks with a step size of 0.75 µm.

### Image quantification

MN were defined as DAPI positive objects near a nucleus that were distinct from the nuclei (not blebs) and lack a teardrop shape (chromatin bridge fragments). Intact MN were defined as those where the H3K27ac mean intensity that was equivalent to that of the nucleus over all or some part of its area or as those with an LBR intensity equivalent to or lower than that of the nucleus. Ruptured MN were defined as those where the H3K27ac mean intensity was decreased by >60% compared to the nucleus or as those with an LBR intensity higher than that of the main nucleus. Chromosome number was defined as the number of centromere foci labeled by CREST. A positive FISH signal was defined as a focus twice the background signal that partially co-localized with a centromere. Projected MN area was calculated from maximum intensity projections by selecting the DAPI channel object using thresholding and either the analyze particles function or the magic wand function in FIJI.

Mean protein intensity quantifications were performed on intact MN and nuclei. Prior to measurement, images were background subtracted using a rolling ball, radius 50 px. A single in-focus confocal section was identified for both the MN and nucleus, and the DAPI or H3K27ac channel was used to define the ROI and measure channel intensity in FIJI. For quantification of sum intensities, ROIs were defined on z-stack sum projections. Ran and RCC1 sum intensities were quantified from z-stacks using Imaris ×64 8.4.2 (Bitplane). An ROI was defined for each MN and nucleus by creating a surface using the DAPI or H3K27ac channels and the mean intensity, total intensity, and volume for each surface were recorded.

H3K29me2 localization in MN was measured on a single in focus section. The MN ROI was generated on the H3K9me2 channel, then divided into concentric shells 1 px wide. The mean fluorescence intensity for each shell was quantified and the number of shells was normalized from 0-1 to compare different sized MN on the same scale.

Nup and CRM1 protein density was quantified in Imaris by first defining an ROI around each MN and nucleus using the contour tool then using the spots tool, with a XY spot diameter set to 0.22 μM, to identify Nup foci. The threshold was adjusted for each image to capture every Nup focus in the nucleus while limiting spots in the cytoplasm. The same threshold was used for each MN nucleus pair. The surface area (µm^2^) was calculated in Imaris from the DAPI channel using the surface tool and Nup density calculated by dividing the total number of foci by the surface area.

Nuclear lamina gaps in STED images were quantified using a previously developed MATLAB pipeline that uses skeletonization and intensity thresholding to define gaps^12^. Nuclear lamina gaps in confocal deconvolved images were quantified using a derivation of this app that allows for non-STED inputs^119^.

### Mutational Signature Analysis

Single nucleotide variant (SNV) data from 50 cancer cohorts from Pan-Cancer Analysis of Whole Genomes (PCAWG) Consortium was downloaded ^120^. Annotations for chromothripsis events matched to coded patient ID i.e. ICGC donor ID from ^58^, were also downloaded leading to a total of 2987 chromothripsis events across 1019 patients across 50 cancer cohorts. Chromothripsis events were previously annotated for: ‘High confidence’, ‘Low confidence’, ‘Linked to high confidence’, and ‘Linked to low confidence’. Only ‘high confidence’ annotated chromothripsis events were selected for analysis and compared to those with annotations of ‘no’ chromothripsis events. Additionally, chromothripsis events annotated as ‘With other complex events’ were removed from the dataset to remove any confounding factors due to other complex events.

Chromosome were categorized into the following categories based on the measured and predicted timing of rupture of MN containing the respective chromosomes^12^:

- Early rupture: ‘13’, ‘18’, ‘21’
- Late rupture: ‘1’,’2’,’3’,’4’,’5’,’6’,’17’,’19’,’X’
- Mid rupture: ‘7’,’8’,’9’,’10’,’11’,’12’,’14’,’15’,’16’,’20’,’22’

The filtered dataset resulted in the following distribution of high confidence chromothripsis events without any other complex events:

| Early Rupture | Mid rupture | Late Rupture |
| --- | --- | --- |
| 37 | 128 | 166 |

Sigprofiler ^121^ was used to assign COSMIC signatures version 3.4 to the SNVs. SNVs in each patient were assigned one of six categories, which were a combination of chromothripsis status of ‘yes’ or ‘no’ and chromosome status of ‘early’, ‘mid’ or ‘late’ rupture. SNVs in each category of the sample were assigned to mutational signatures, which were then normalized by the total number of SNVs in the sample category. The normalized proportion of SNVs assigned to APOBEC signature (SBS 2 and SBS 13) or the PolE signature (SBS10a and b) were averaged across early, mid and late rupture chromosomes that were chromothripsis positive. Code to process the data can be found on a public repository here: https://github.com/manasvitavashisth/MutationalSignatureAnalysis

### Statistics

All statistic tests were conducted using R (version 4.0.0) or Prism. For all tests, *P*-values greater than 0.05 were considered statistically significant. For all data comprising three or more groups of observations, a family test (i.e., chi-squared for categorical data, ANOVA test for continuous data) was performed first in R using the R function ‘aov’. Only data where the family test was significant were further analyzed by multiple comparison testing. For statistical analysis of categorical data between 2 groups, *P*-values were calculated using Barnard’s exact test in R using the ‘Barnard’ package (version 3.4.1). For statistical analysis of the distribution of quantitative data between 2 groups where one or more groups deviated from normality, *P*-values were calculated in R with Wilcoxon rank sum test (base R). For statistical analysis to determine whether a dataset differed from a defined number, *i.e.* 1, *P*-values were calculated using a one sample Wilcoxon rank sum test in R (base R). Spearman’s rank correlation coefficient was used to assess monotonic relationships for two variables with non-normal distribution with R or Prism. For analysis of chromothripsis mutational signatures, the Tukey Honest Significant Differences test was used to used ascertain statistically significant differences in the increase in APOBEC and PolE signature proportion in the early and mid rupture chromosomes with chromothripsis, respectively, in R (base R). Shorthand p-values are as follows:

ns: p-value >= 0.05

*: p-value < 0.05

**: p-value < 0.01

***: p-value < 0.001

****: p-value < 0.0001

## Acknowledgments

E.M.H was supported by the National Institutes of Health grant R35GM124766 and a Rita Allen Foundation Scholars Award. M.G.Z was supported by a National Institutes of Health training grant (T32 GM007270). A.E.M. was supported by a National Institutes of Health training grant (T32CA009657). G.H. was supported by a National Institutes of Health grant DP2CA280624. This work was supported by the Cellular Imaging and Flow Cytometry Shared Resources of the Fred Hutch/University of Washington Cancer Consortium (P30 CA015704).

## Author Contributions

MG Zych: conceptualization, data curation, formal analysis, funding acquisition, investigation, methodology, project administration, validation, visualization, writing – original draft, review and editing

M Contreras: data curation, formal analysis

M Vashisth: formal analysis

A Mammel: data curation

G Ha: project administration, supervision

EM Hatch: conceptualization, formal analysis, funding acquisition, investigation, methodology, project administration, resources, supervision, visualization, writing – original draft, review & editing

## Figure legends

**Supplemental table 1. Cancer cohorts for chromothripsis events**

For figure 6, the distribution of chromothripsis events across the cancer cohorts from the PCAWG dataset that were included in our analysis.

**Supplemental table 2. Chromothripsis event data used in signature analysis**

For figure 6, raw data from PCAWG dataset for the chromothripsis events that were used in our analysis to identify DNA damage signatures.

**Supplementary Figure 1.**
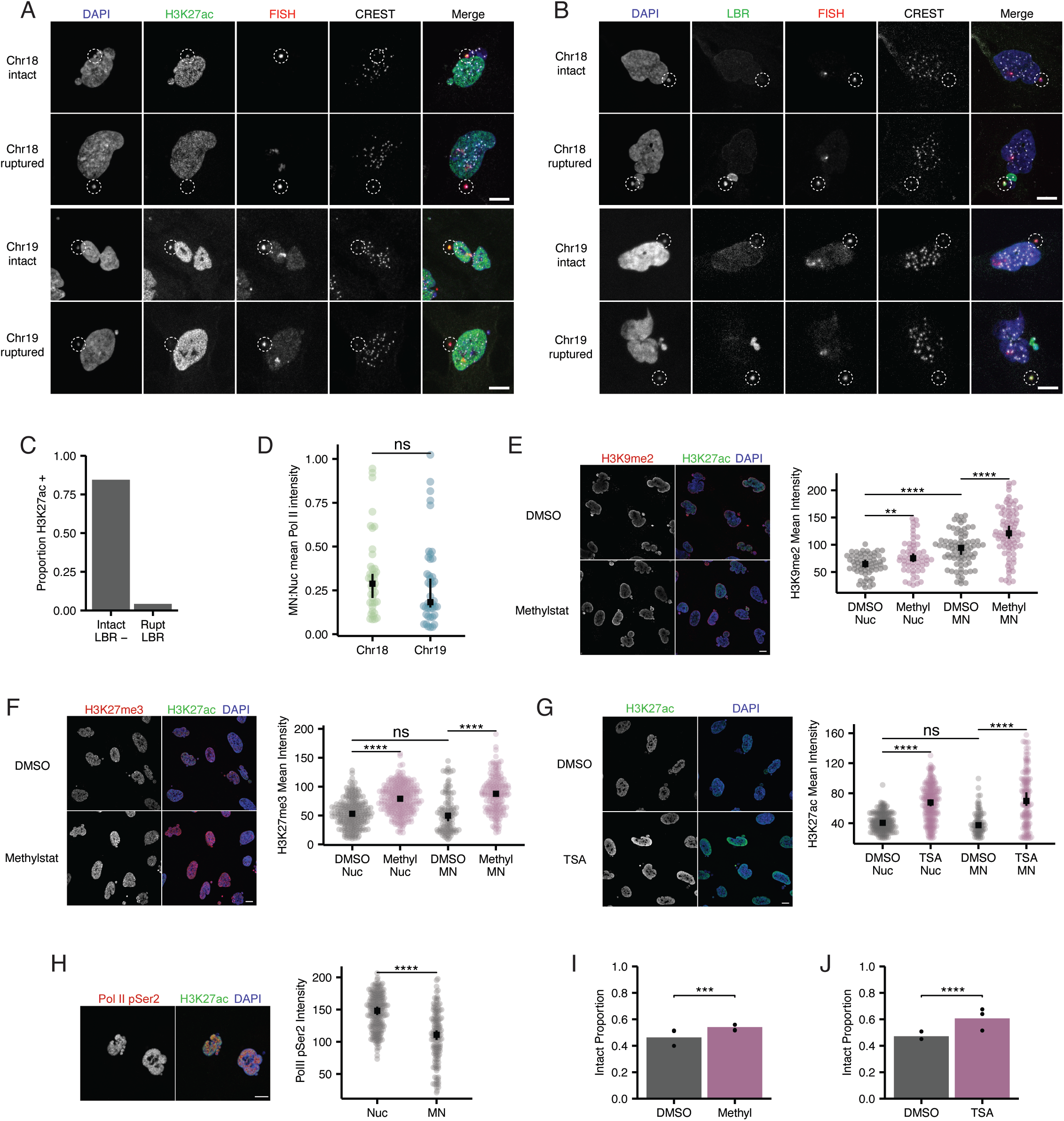
Histone modifications are altered in MN and impacted by drug treatments. **A,B.** Maximum intensity projection images of **A.** intact (H3K27ac+) and ruptured (H3K27ac-) MN and **B.** intact (LBR-) and ruptured (LBR+) MN in RPE1 cells containing a single copy (1 CREST focus) of chr 18 or 19. Scale bars for A and B = 10um. **C.** Quantification of H3K27ac and LBR colocalization in RPE1 cells 84.5% LBR-MN were of H3K27ac+ and 4.3% of LBR+ MN were of H3K27ac+. N=3, n=620. **D.** MN to nuclear ratio of mean RNA PolII pSer2 intensity for intact single chromosome chr 18 and 19 MN in RPE1 cells 24hrs post BAY addition. Wilcox ran sum test, N=3, n=(16, 15). **E,F**. Example images of H3K9me2 and H3K27me3 staining in nuclei and MN from RPE1 cells with single sections shown. Quantification of H3K9me2 and H3K27me3 intensity in nuclei and intact (H3K27ac+) MN in cell treatment with DMSO and 5uM methylstat. Scale bar = 10um. **E.** H3K9me2 Wilcoxon ran sum test, N=3, n=(62,58,82,86) **F.** H3K27me3 Wilcoxon ran sum test, N=3, n=(231, 269, 109, 177). **G.** Example images of H3K27ac staining nuclei and MN from RPE1 cells with single sections shown. Quantification of H3K27ac in nuclei and micronuclei in cells treated with DMSO and 100nM TSA . Scale bar = 10um. Wilcoxon ran sum test. N=3, n=(224, 252, 93, 149). **H.** Example images of Pol II pSer2 staining in nuclei and MN from RPE1 cells with single sections shown. Quantification of Pol II pSer2 intensity in nuclei and intact (H3K27ac+) MN. Wilcoxon ran sum test. N=3, n=(210, 136). **I.** Intact proportion of bulk MN 24hr post BAY determined by H3K27ac classification in cells treated with DMSO and 5uM methylstat. Barnards test, N=3, n=(477,432, 386). **J.** Intact proportion of bulk MN 24hr post BAY determined by LBR classification in cells treated with DMSO and 100nM TSA. Barnards test, N=3, n=(462, 560). For all graphs, ns p>0.05, * p<0.05, ** p<0.01, *** p<0.001, **** p<0.0001.

**Supplementary Figure 2.**
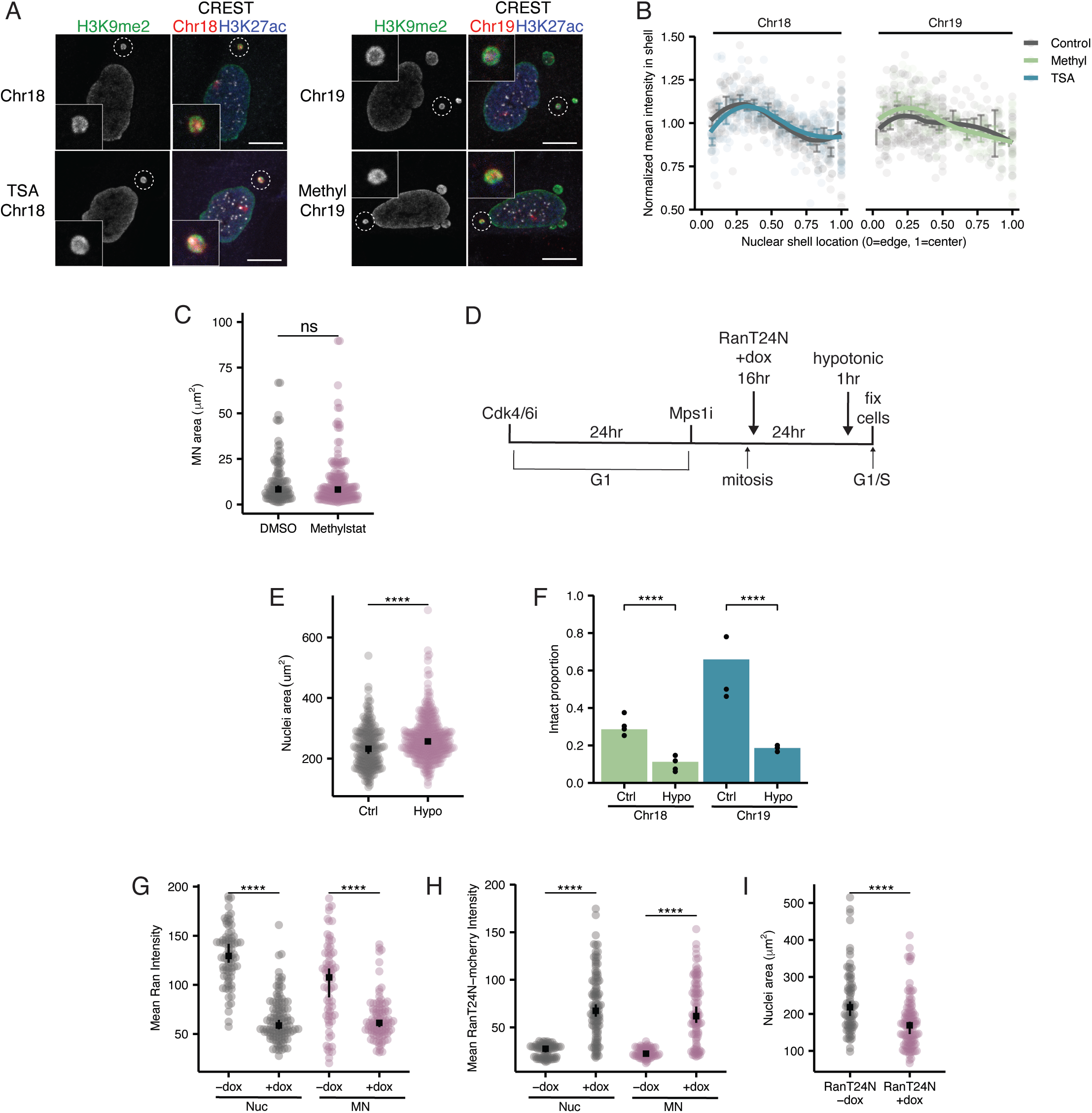
Lamina gaps and instability in MN correlate with growth not LADs. **A.** Example images of H3K9me2 staining in single chromomsome 18 and 19 RPE1 cell MN treated with DMSO, 5uM methylstat, or 100nM TSA with single sections shown. Scale bar = 10um. **B.** Chr 18 and 19 MN from A were segmented into 1-pixel thick concentric shells and the mean H3K9me2 intensity was quantified in each shell. The intensity of each shell is normalized to mean intensity of the whole MN and is shown with the standard error. N=3, chr 18 n=(30, 34), chr 19 n=(29, 30). **C.** Max projected area for bulk intact MN in RPE1 cells treated with DMSO and 5uM methystat for 4 hours. N=3, n=(109, 177). **D.** Experimental outline of 1hr hypotonic treatment and 16hr doxycycline treatment for RanT24N-mcherry induction. **E.** Maximum projected area of nuclei in RPE1 control cells and after hypotonic swelling in media diluted 1:2 in H2O for 1 hour. Wilcoxon rank sum test, N= 3, n=(83,135). **F.** MN stability for single chromosome 18 or 19 MN in RPE1 cells 24hrs after BAY for control and hypotonic treated cells. Barnards test, N=3, n=(224, 169, 85, 70). **G.** Quantification of Ran staining in nuclei and intact MN in RPE1 TetOn Tre-RanT24N-mcherry cells treated with 0ng/ul and 100ng/ul dox. Wilcoxon rank sum, N=3, n=(79,113,72,97). **H.** Quantification of RanT24N-mcherry staining in nuclei and intact MN in RPE1 TetOn Tre-RanT24N-mcherry cells treated with 0ng/ul and 100ng/ul dox. Wilcoxon rank sum, N=3, n=(79,113,72,97). **I.** Max projected area of nuclei in RPE1 TetOn TRE-RanT24N-mcherry cells treated with 0ng/ul or 100ng/ul dox for 16hrs. Wilcoxon rank sum test, N=3, n=(79,113). For all graphs, ns p>0.05, * p<0.05, ** p<0.01, *** p<0.001, **** p<0.0001.

**Supplementary figure 3:**
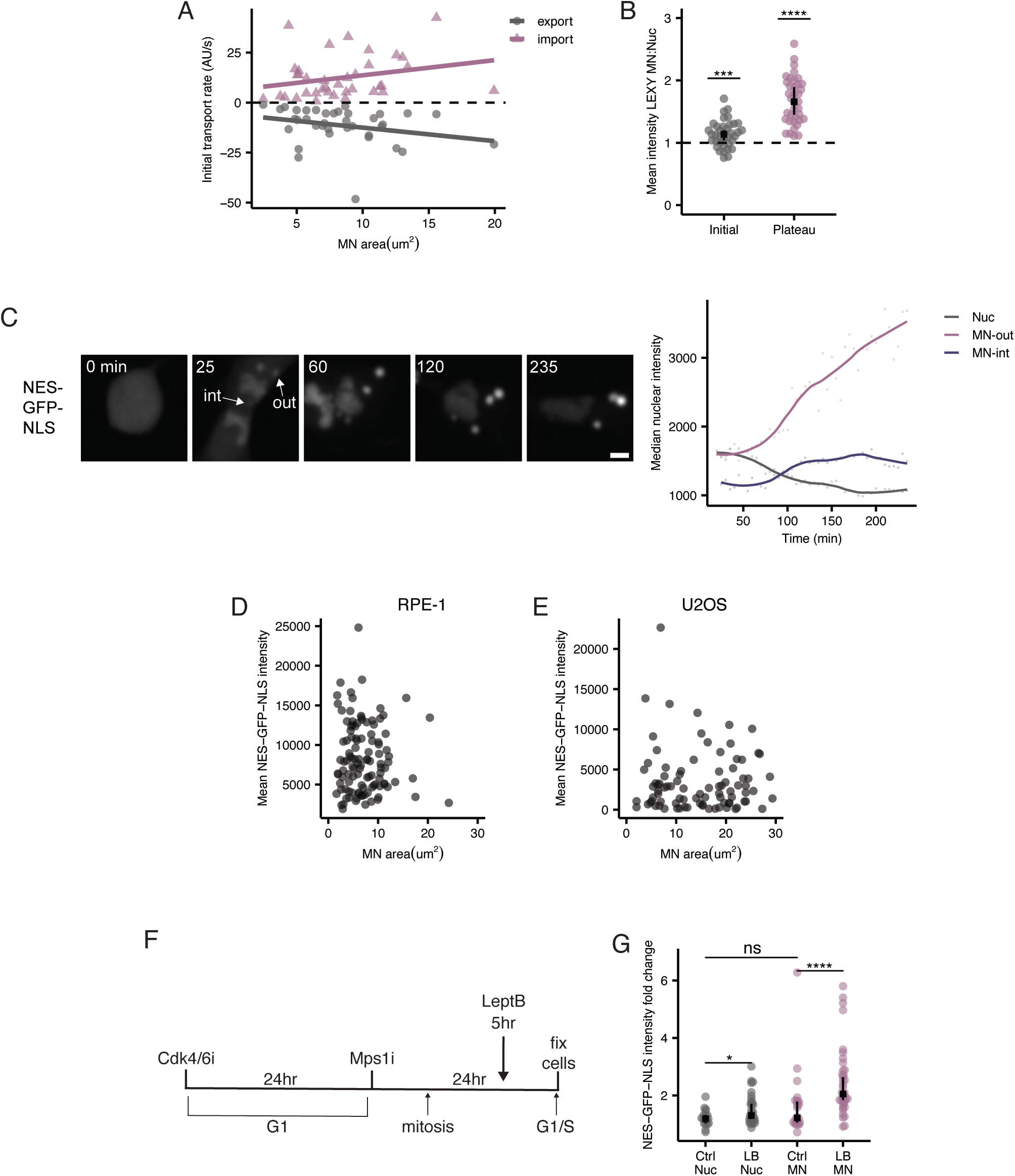
MN transport rates are conserved and not correlated with MN area. **A.** Nuclear import and export rates quantified from LEXY-mCherry intensity changes are not correlated with MN area. **B.** LEXY-mCherry intensity quantified in MN and nuclei at the start of imaging in the absence of UV (initial) and after 45 min of UV exposure to induce export (plateau). MN were compared to their corresponding nuclei. One-sample Wilcoxon rank sum test, N=3, n=(34). **C.** Live imaging of RPE1 NES-GFP-NLS cells post mitosis. Time 0 starts at mitotic entry and is shown in minutes. Scale bar = 10um. Quantification of the median NES-GFP-NLS intensity in nuclei and MN over time is shown for MN in the interior of the spindle (MN int) and MN outside the spindle (Mn-out). **D.** NES-GFP-NLS intensity levels quantified in RPE1 cell MN are not correlated with MN area. Spearman correlation R= -0.03, p=0.75. **E.** NES-GFP-NLS intensity levels quantified in U2OS cell MN are not correlated with MN area. Spearman correlation R= 0.017, p=0.94. **F.** Experimental outline of 5hr leptomycin B treatment. **G.** The intensity of NES-GFP-NLS for nuclei and intact MN measured from live imaging of RPE1 cells expressing NES-GFP-NLS at the start of imaging and 5hrs after 20ng/ul leptB LB addition. The fold change of NES-GFP-NLS intensity over 5hrs was calculated. Wilcoxon rank sum test, N=1, Nuc n=(19, 31), MN n=(24, 39). For all graphs, ns p>0.05, * p<0.05, ** p<0.01, *** p<0.001, **** p<0.0001.

**Supplementary figure 4:**
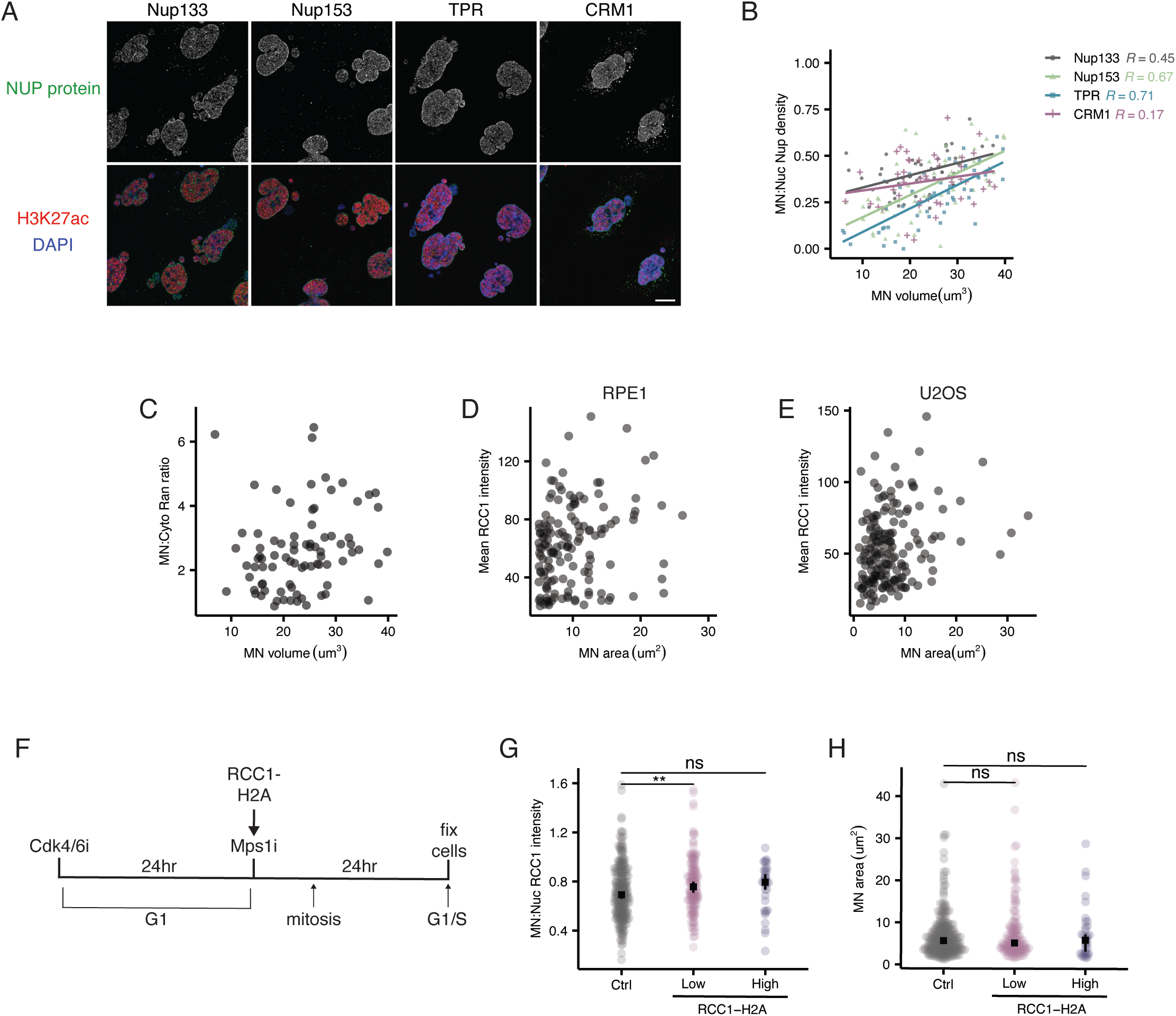
RCC1 levels more than NUP density correlate with export defects in MN. **A.** Maximum projections of the top surface of nuclei and MN in RPE1 cells stained for Nup133, Nup153, TPR, and CRM1. Scale bar =10um. **B.** Correlation between NUP density and MN volume for quantification in 4A. R= Spearman correlation coefficient. Nup133 p<0.01, Nup153 p<0.0001, TPR p<0.0001, CRM1 p>0.05. **C.** Ran intensity levels quantified in RPE1 cell MN are not correlated with MN area. Spearman coefficient R= 0.26, p=0.0017. **D.** RCC1 intensity levels quantified in RPE1 cell MN are not correlated with MN area. Spearman coefficient R= 0.24, p=0.0033. **E.** RCC1 intensity levels in U2OS cell MN are not correlate with MN area. Spearman coefficient R= 0.1, p=0.41. **F.** Experimental outline of RCC1-H2A expression for 24hrs. **G.** MN:Nuc ratio of RCC1 intensity in control and RCC1-H2A expressing RPE1 cells**. H.** Area of MN in control and RCC1-H2A expressing RPE1 cells show no change in MN size with RCC1 overexpression. **G,H.** Wilcoxon rank sum test, N=4, n=(270, 149, 36). For all graphs, ns p>0.05, * p<0.05, ** p<0.01, *** p<0.001, **** p<0.0001.

**Supplementary figure 5:**
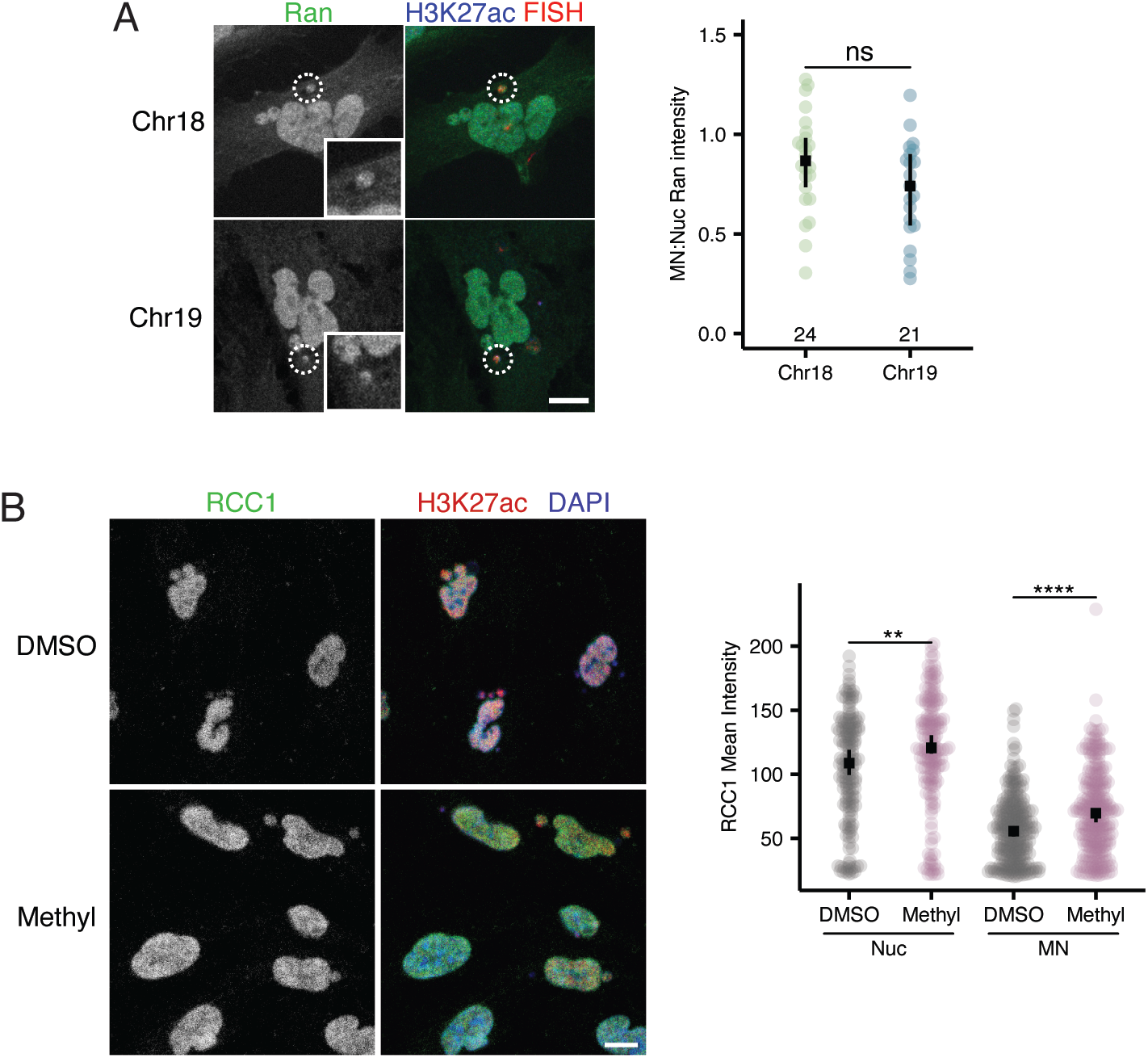
Transport factors in MN are correlated with histone methylation. **A.** Maximum projections of Ran in intact single chromosome 18 and 19 MN in RPE1 cells 24hrs post BAY addition. Wilcoxon rank sum test, N=3, n=(24, 21). **B.** Images of RCC1 in RPE1 cells treated with DMSO and 5um methylstat. Single sections shown. Quantification of RCC1 intensity in nuclei and intact MN. Wilcoxon ran sum test, N=3, n=(168, 177, 268, 276). For all graphs, ns p>0.05, * p<0.05, ** p<0.01, *** p<0.001, **** p<0.0001.

**Supplemental figure 6.**
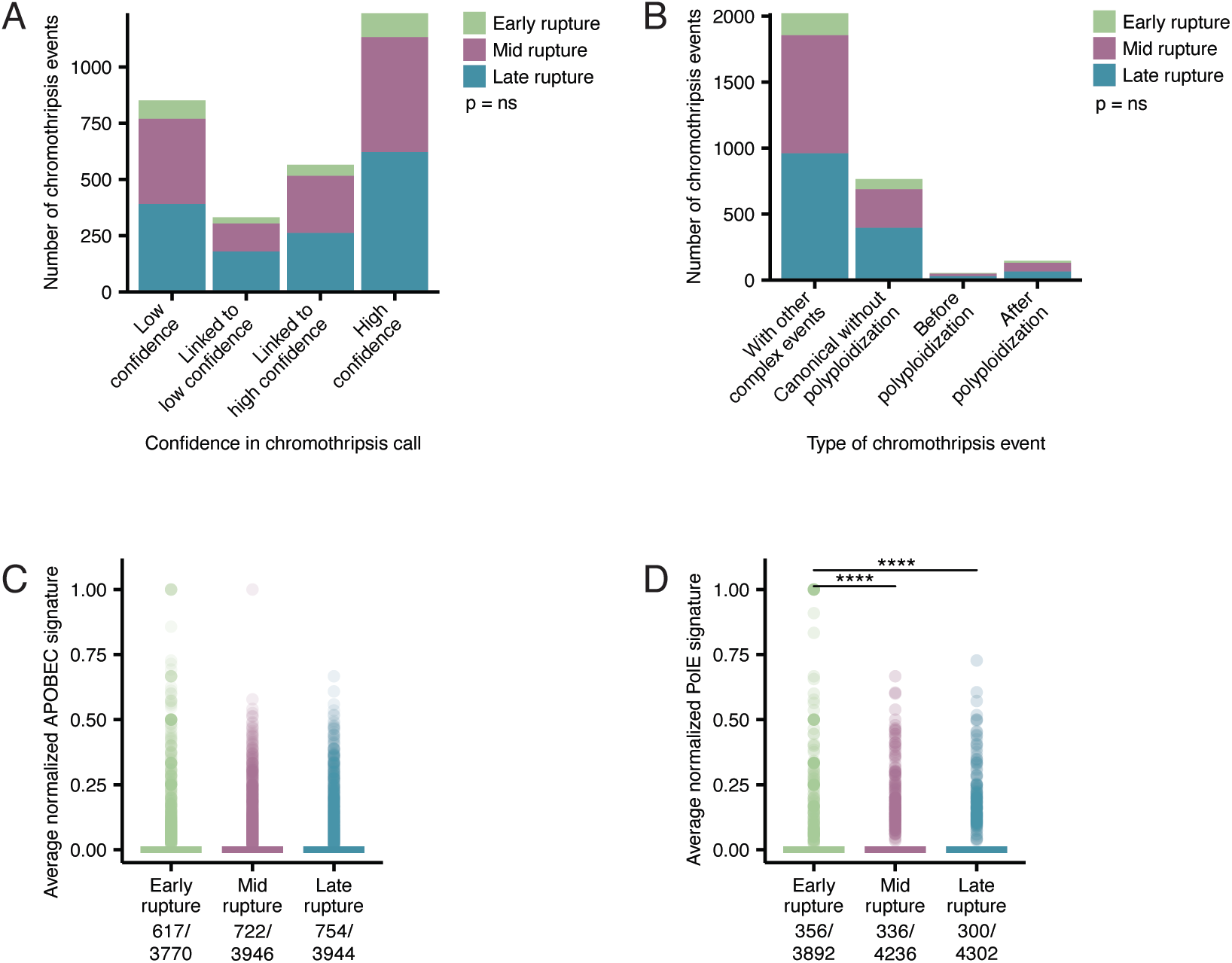
Controls for mutational signature enrichment in chromothripsis events. **A,B.** Rupture timing classes are equally distributed between different chromothripsis annotations and chromothripsis event annotations. **C.** APOBEC mutational signature prevalence is not increased in early rupture classes annotated as “no” chromothripsis. p= ns, ANOVA. Number of non-zero datapoints out of the total are indicated under each group. n= 3770, 3946, 3944. **D.** PolE mutational signature is not enriched in mid rupture classes annotated as “no” chromothripsis. p= <0.0001, ANOVA. Number of non-zero datapoints out of the total are indicated under each group. n= 3892, 4236, 4302. For all graphs, ns p>0.05, * p<0.05, ** p<0.01, *** p<0.001, **** p<0.0001.

